# The transcriptomic landscape of the photoperiodic stress response in *Arabidopsis thaliana* resembles the response to pathogen infection

**DOI:** 10.1101/2021.04.13.439491

**Authors:** Anne Cortleven, Venja M. Roeber, Manuel Frank, Jonas Bertels, Vivien Lortzing, Gerrit Beemster, Thomas Schmülling

## Abstract

Plants are exposed to regular diurnal rhythms of light and dark. Changes in the photoperiod by the prolongation of the light period cause photoperiod stress in short day-adapted *Arabidopsis thaliana*. Here we report on the transcriptional response to photoperiod stress of wild-type *A. thaliana* and photoperiod stress-sensitive cytokinin signalling and clock mutants. Transcriptomic changes induced by photoperiod stress included numerous changes in reactive oxygen (ROS)-related transcripts and showed a strong overlap with changes occurring in response to ozone stress and pathogen attack, which have in common the induction of an apoplastic oxidative burst. A core set of photoperiod stress-responsive genes has been identified, including salicylic acid (SA) biosynthesis and signalling genes. Genetic analysis revealed a central role for NPR1 in the photoperiod stress response as *npr1-1* mutants were stress-insensitive. Photoperiod stress treatment led to a strong increase in camalexin levels which is consistent with shared photoperiod stress and pathogen response pathways. Photoperiod stress induced resistance of *Arabidopsis* plants to a subsequent infection by *Pseudomonas syringae* cv. *tomato* DC3000 indicating priming of the defence response. Together, photoperiod stress causes transcriptional reprogramming resembling plant pathogen defence responses and induces systemic acquired resistance in the absence of a pathogen.

**One sentence summary:** Photoperiod stress results in significant dynamic transcriptomic changes related to oxidative stress similar to those caused by pathogen attack and primes the defence response against a subsequent pathogen infection.

## INTRODUCTION

The photoperiod is the duration of the light period during a daily day-night cycle of 24 h. As the earth turns around its own axis, the daily change of day and night results in the adaptation of life processes to this rhythm. As a consequence, numerous developmental processes are controlled by the photoperiod (Jackson, 2009). Recently, it has been described that changes of the photoperiod, in particular a prolongation of the light period, provokes a stress response in *Arabidopssi thaliana*. This newly identified form of abiotic stress was named photoperiod stress - originally circadian stress - (Nitschke et al., 2016; Nitschke, Cortleven, & Schmulling, 2017) and was first detected in cytokinin (CK)-deficient *Arabidopsis* plants as these plants are particularly stress-sensitive. The photoperiod stress response starts during the night following a prolonged light period with a strong induction of stress marker genes such as *ZAT12* (*ZINC FINGER of ARABIDOPSSI THALIANA2*) and *BAP1* (*BON ASSOCIATED PROTEIN1*), an increase in oxidative stress and jasmonic acid (JA) level. The next day, photosynthetic efficiency is strongly reduced and eventually programmed cell death follows. A weaker molecular response during the night following the extended light period was also detected in wild type (WT) (Abuelsoud, Cortleven, & Schmülling, 2020; Nitschke et al., 2016). The nightly increase in oxidative stress coincides with a strong decrease in ascorbic acid (ASC) redox state and the formation of peroxides (including H_2_O_2_), which is associated with a strong increase of *PEROXIDASE* (*PRX*) gene expression, enhanced PRX activity and reduced catalase activity (Abuelsoud et al., 2020). CK, especially root-derived *trans*-zeatin forms (Frank et al., 2020), have a protective function acting through the ARABIDOPSIS HISTIDINE KINASE 3 (AHK3) receptor and the transcriptional regulators ARABIDOPSIS RESPONSE REGULATORS ARR2, ARR10 and ARR12. Also, certain clock mutants of both the morning and evening loops (e.g. *cca1 lhy, elf3*) are photoperiod stress-sensitive (Nitschke et al., 2016). Common to stress-sensitive clock mutants and CK-deficient plants was a lowered expression or impaired functioning of CIRCADIAN CLOCK ASSOCIATED1 (CCA1) and LONG HYPOCOTYL (LHY), two key regulators of the circadian clock, which indicates that a functional clock is essential to cope with stress caused by altered light-dark rhythms (Nitschke et al., 2016). An important function of the circadian clock is to anticipate the daily light-dark rhythm and matching the circadian clock with the environment is crucial for the regulation of numerous biological processes including the activity of plant hormones, the formation of and response to reactive oxygen species (ROS) and the responses to plant pathogens (Carmela, Ewers, & Weinig, 2018; Covington, Maloof, Straume, Kay, & Harmer, 2008; Harmer, 2009; Karapetyan & Dong, 2018; Roden & Ingle, 2009; Seo & Mas, 2015).

Plant defense responses to pathogens involve a multilayered strategy including the primary innate immunity and a host-specific secondary immune response (Chisholm, Coaker, Day, & Staskawicz, 2006; de Wit, 2007). Pathogen-associated molecular patterns (PAMPs) are detected by pattern recognition receptors resulting in PAMP-triggered immunity (PTI). This primary innate immunity involves the induction of pathogen-responsive genes, ROS production or alterations in hormone signaling pathways involving salicylic acid (SA) and JA. Certain pathogens produce effector proteins that are encoded by avirulence genes to circumvent the PTI. These pathogen-derived effectors can be counteracted by plant resistance proteins encoded by *R* genes resulting in effector-triggered immunity (ETI). Both PTI and ETI lead to similar plant responses and have comparable signaling pathways involving ENHANCED DISEASE SUSCEPTIBILITY1 (EDS1) and PHYTOALEXIN4 (PAD4) which promote *ICS1* (*ISOCHORISMATE SYNTHASE1*) expression and thus SA accumulation (Cui et al., 2017). Increased cellular SA levels activate several signaling cascades in which NON-EXPRESSOR OF PATHOGEN-RELATED GENE1 (NPR1) has a central role as master regulator. NPR1 interacts with transcription factors to induce the production of antimicrobial peptides such as pathogenesis-related (PR) proteins (Backer, Naidoo, & van den Berg, 2019; Pajerowska-Mukhtar, Emerine, & Mukhtar, 2013; Qi et al., 2018). SA accumulation induces biosynthesis of camalexin, which is the major phytoalexin formed during biotic stress responses (Glawischnig, 2007). EDS1 and PAD4 also regulate a SA-independent immunity pathway in which FLAVINDEPENDENT MONOOXYGENASE1 (FMO1) acts as an inducer of systemic acquired resistance (SAR) (Bartsch et al., 2006; Hartmann et al., 2018) by regulating the production of N-hydoxypipecolic acid (NHP).

Light is crucial for the activation of full resistance responses in plant-pathogen interactions, for both the induction of local defense responses as well as for the transcriptional regulation required for SAR (for review: Ballaré (2014); Roeber, Bajaj, Rohde, Schmulling, and Cortleven (2020)). Several studies have shown that in particular the length of the light period determines the strength of the plant immune response in *Arabidopssi thaliana* (Cecchini et al., 2002; Griebel & Zeier, 2008). The length of the photoperiod also plays an important role in other stress responses, including the response to cold (C. M. Lee & Thomashow, 2012) and heat (Dickinson et al., 2018; Han, Park, & Park, 2019).

The experience of stress can prepare plants to react more rapidly or even stronger to recurrent stress of the same or different nature, a process that is called priming (Hilker & Schmulling, 2019; Hilker et al., 2016). The activation of defense mechanisms in response to an initial pathogen attack resulting in SAR is one of the best-characterized forms of priming (Conrath, Thulke, Katz, Schwindling, & Kohler, 2001). Other well-known examples of priming are cold acclimation (Thomashow, 2010) and thermotolerance (Song, Jiang, Zhao, & Hou, 2012) which enables plants to survive normally lethal temperatures after previous exposure to non-lethal low or high temperatures, respectively. Also light may prime plants to better resist high temperatures, indicating that the priming stress may be different from the second, triggering stress (Han et al., 2019).

In this study, we have investigated transcriptional changes occurring in response to photoperiod stress. We have compared the transcriptomic landscape in WT plants and in two photoperiod stress-sensitive mutants, *ahk2 ahk3* and *cca1 lhy* during the course of the night following a prolonged light period (PLP). Photoperiod stress results in strong changes of transcript abundance, which show a distinct time-dependent profile. Among the differentially expressed genes (DEGs) responding to photoperiod stress are many ROS-responsive genes. Globally, the transcriptional changes caused by photoperiod stress resemble those induced by ozone stress and pathogen attack including deregulated expression of genes related to SA biosynthesis and signaling. NPR1, an important regulator of the plant pathogen response is required for the response to photoperiod stress. An enhanced resistance of *Arabidopsis* to pathogen infection after a preceding photoperiod stress exposure is consistent with the similarities between the two stresses and indicates priming by photoperiod stress.

## RESULTS

### Transcriptional dynamics of the photoperiod stress response

Changes of transcript abundance in response to photoperiod stress were analyzed in five-weeks-old short day-grown plants (WT, *ahk2 ahk3, cca1lhy*) during the night following a light period that was prolonged by 24 h. We used this treatment causing a strong stress response (Nitschke et al., 2016) but it should be noted that shorter prolongation of the photoperiod in the range of few hours also causes a significant stress response (Abuelsoud et al., 2020). Samples for RNA-seq analysis were harvested at different time points (0 h, 4 h, 6 h and 12 h) following the prolonged light treatment (Fig. 1A). These time points reflect the timing of stress responses occurring during the night with stress marker gene activation starting around 4 h and visible phenotypical consequences (flabby leaves) appearing about 8 h after the start of the night. Thus, the time points allow monitoring of early transcriptional changes occurring before the onset of the stress symptoms. A scheme of the experimental setup is shown in Fig. 1A.

**Figure 1.**
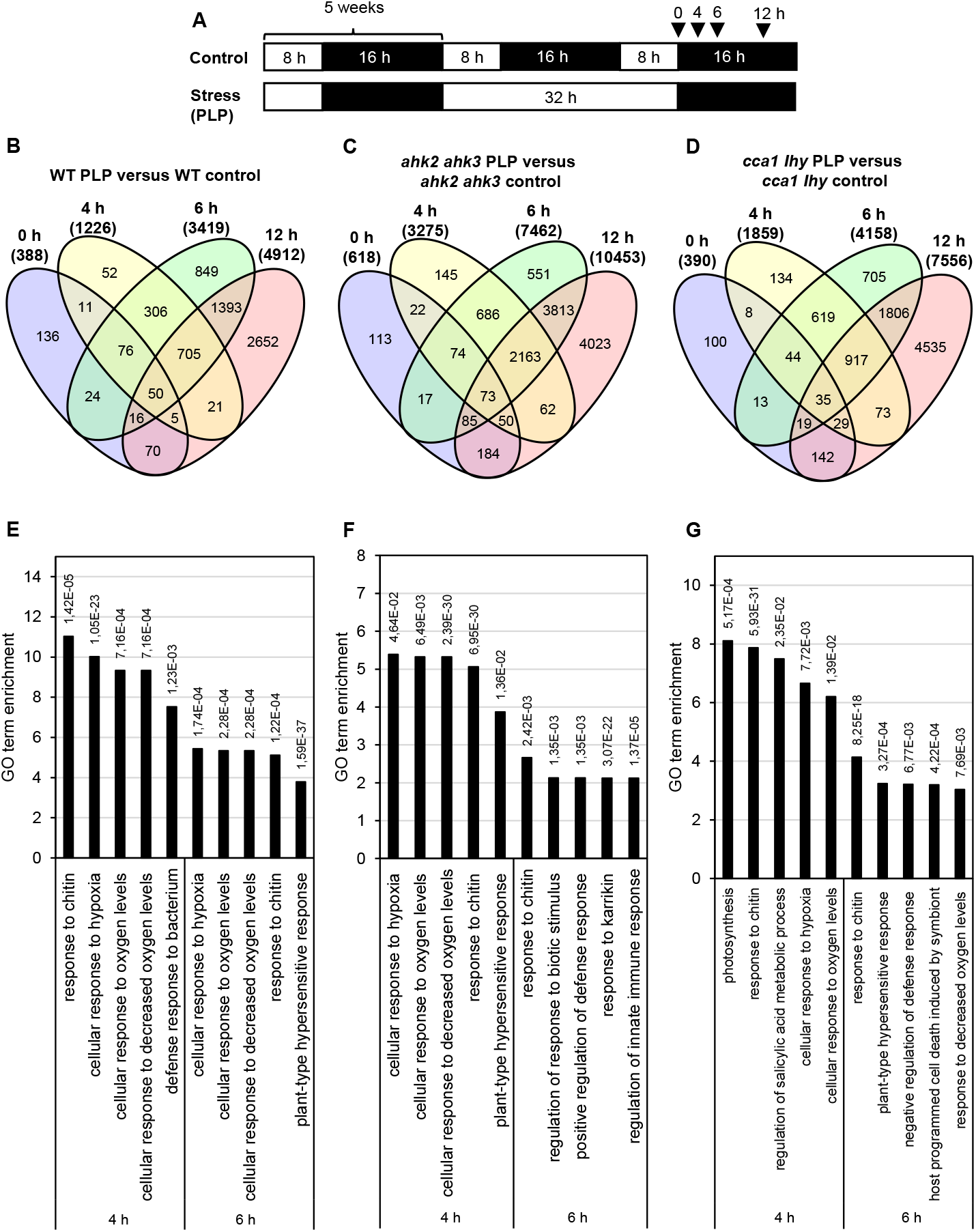
Significantly regulated genes in WT, *ahk2 ahk3* and *cca1 lhy* after photoperiod stress. (A) Experimental setup used in this study. 5-weeks-old short-day grown plants were exposed to a prolonged light period (PLP) of 32 h followed by a normal short-day night. White; light period; black, dark period. Arrows indicate sampling time points for RNA analysis. (B - D) Venn diagrams showing the overlap of DEGs at different time points for WT (B), *ahk2 ahk3* (C) and *cca1 lhy* (D). Numbers in brackets indicate the total number of DEGs (|fold-change| = 2; Bonferroni-corrected p-value ≤ 0.05) in PLP-treated plants compared with control plants at the different time points. (E - G) Top 5 GO enrichment for time point 4 h and 6 h for WT (E), *ahk2 ahk3* (F) and *cca1 lhy* (G). The complete list of GO enrichment terms can be found in Supplemental Table S1. An overview of the gene regulation for the comparisons between PLP and control treatments for WT, *ahk2 ahk3* and *cca1 lhy* is shown for all time points in Supplemental Data 2. A core-set of photoperiod stress-responsive genes is listed in Supplemental Table S2 and the top 20 most highly regulated genes at each time point are listed in Supplemental Tables S4-S9. PLP; prolonged light period.

Principal component analysis (PCA) of the DEGs showed that control samples cluster together (red circle; Supplemental Fig. S1A) indicating that changes in gene expression caused only by the genotype are much smaller than those caused by the treatment. At the end of the PLP, the 0 h time point belongs to the cluster of the control samples suggesting that the photoperiod stress response has not yet started. The 4 h and 6 h time point cluster separate from control samples and a division among the different genotypes is visible. Especially at the 12 h time point, a strong separation of the genotypes (blue circles; Supplemental Fig. S1B) is evident. At this time point, the *ahk2 ahk3* and *cca1 lhy* samples are clearly divergent from WT, which fits to the stronger photoperiod stress phenotype of these mutants (Nitschke et al., 2016). Statistical analysis of the RNA-seq data set was performed with DEseq2 considering three different parameters (time, genotype and treatment) and corrected with Bonferroni. There are 10278 genotype-dependent genes, 17465 genes are significantly regulated by time and 33208 genes are treatment-dependent considering all different time points and genotypes. This indicates that the effect of the treatment is larger than the time- or genotype-effect. The genotype-dependent differentially expressed genes were analyzed in detail by cluster and GO term analysis. A Quality Treshold (QT) clustering was performed and revealed 21 different clusters (Supplemental Fig. S2). 52 % of all DEGs are found in cluster 1 and 2 which showed an upregulation (cluster 1) or a downregulation (cluster 2) during the night following the photoperiod stress treatment. This regulation has a higher amplitude in the stress-sensitive mutants (Supplemental Fig. S1B-C). According to GO term analysis, mostly genes involved in photosynthesis- or chloroplast-related processes belong to cluster 1 and genes involved in autophagy, responses to endoplasmatic reticulum stress and Golgi vesicle transport belong to cluster 2 (Supplemental Fig. S1D-E).

To get an insight in the DEGs following photoperiod stress, pairwise comparisons between photoperiod stress-treated and control samples were made for each genotype and time point. As thresholds for DEGs, a fold change ≥ |2| and a p-value ≤ 0,05 (Bonferroni-corrected) were chosen. An overview of all gene identifiers of significant DEGs in WT, *ahk2 ahk3* and *cca1 lhy* after photoperiod stress expressed relative to control treatment pro time point can be found in Supplemental Data 1. In Supplemental Data 2, fold change and significance levels for the comparison between prolonged light period (PLP) and control treatment for WT, *ahk2 ahk3* and *cca1 lhy* at time points 0 h, 4 h, 6 h and 12 h are shown.

An overview of the number of regulated genes over time is shown in Figure 1B-D. In all genotypes, the number of DEGs increases over time. For instance, in WT there are 388 DEGs at time point 0 h, 1226 DEGs at time point 4 h, 3419 DEGs at time point 6 h and 4912 DEGs at time point 12 h. A large number of the early regulated genes show also an increased steady state level at later time points. Among the DEGs increases the number of TF genes over time suggesting that the stress response involves transcriptional cascades. In the stress-sensitive mutants, a similar pattern is seen. The number of DEGs increases over time, however the number of regulated genes is considerably higher. The total number of DEGs at time point 12 h is 10453 in *ahk2 ahk3* and 7556 in *cca1 lhy* reflecting their higher sensitivity.

Among the top five GO terms at the 4 h and 6 h are in all genotypes cellular responses to oxygen levels and plant-type hypersensitive responses or positive regulation of defense responses (Figure 1E-G). This indicates that the transcriptomic landscape of photoperiod stress can be associated with oxidative stress. An overview of all GO enrichment terms can be found for all time points in Supplemental Table S1.

The DEGs that are shared between WT, *ahk2 ahk3* and *cca1 lhy* at time points 4 h, 6 h and 12 h resulted in a core set of 388 photoperiod stress-regulated genes, 382 genes were upregulated and 6 genes were downregulated (Table1, Supplemental Table S2). A prolongation of the light period alone is not causative for the photoperiod stress response but it is triggered by the following dark period (Nitschke et al., 2016). Therefore, we have omitted the 0 h time point as the observed changes are genotype-dependent light effects caused by the prolongation of the light period and do not belong to the specific photoperiod stress response occurring during the night. However, DEGs at time point 0 h might be relevant for the perception of photoperiod stress and the development of the initial response. An overview of the significantly genes regulated at time point 0 h is given in Supplemental Table S2. GO term analysis revealed that the core set of photoperiod stress-regulated genes belong mostly to cellular responses to hypoxia or to oxygen levels and defense responses to pathogens. This supported further a strong correlation between photoperiod and oxidative stress (Supplemental Table S1).

**Table 1:** Selection of the core-set of photoperiod stress responsive genes. Fold changes are sorted according to the 4 hours timepoint in WT. Only statistical significant differently regulated genes are shown (Bonferroni < 0.05). FC, fold change. The complete core-set of photoperiod stress responsive genes can be found in Supplemental Table S2.

**upregulated DEGs**
| ATG | WT<br>(log<sub>2</sub> FC) |  |  | *ahk2 ahk3*<br>(log<sub>2</sub> FC) |  |  | *cca1 lhy*<br>(log<sub>2</sub> FC) |  |  | short description |
| --- | --- | --- | --- | --- | --- | --- | --- | --- | --- | --- |
|  | 4 h | 6 h | 12 h | 4 h | 6 h | 12 h | 4 h | 6 h | 12 h |  |
| AT3G46080 | 7,42 | 6,63 | 7,46 | 5,87 | 9,53 | 4,36 | 7,03 | 9,01 | 4,89 | C2H2-type zinc finger family protein |
| AT2G45760 | 5,94 | 7,47 | 5,63 | 6,24 | 8,84 | 4,39 | 8,50 | 8,90 | 4,63 | encodes a protein that is similar to BONZA1-binding protein BAP1 (BAP2) |
| AT4G01360 | 5,91 | 6,79 | 6,74 | 6,14 | 8,90 | 3,87 | 7,28 | 8,23 | 3,91 | encodes a protein related to BYPASS1 (BPS3) |
| AT1G07160 | 5,74 | 6,31 | 5,31 | 5,53 | 6,59 | 3,24 | 6,91 | 8,02 | 3,01 | Protein phosphatase 2C family protein |
| AT4G08555 | 5,55 | 8,49 | 7,87 | 5,45 | 8,54 | 4,12 | 7,02 | 9,66 | 4,65 | hypothetical protein |
| AT1G26380 | 5,52 | 6,97 | 8,19 | 5,22 | 7,44 | 4,43 | 6,74 | 9,00 | 4,28 | FAD-LINKED OXIDOREDUCTASE 1 (FOX1) |
| AT5G66890 | 5,46 | 10,34 | 7,69 | 6,77 | 11,22 | 4,51 | 8,85 | 12,08 | 4,98 | N REQUIREMENT GENE 1.3 (NRG1.3) |
| AT2G32210 | 5,42 | 6,70 | 6,05 | 5,87 | 7,62 | 3,85 | 6,29 | 7,87 | 4,24 | CYSTEINE-RICH TRANSMEMBRANE MODULE 6 (ATHCYSTM6) |
| AT5G64870 | 5,37 | 5,72 | 4,61 | 6,17 | 7,93 | 3,18 | 6,44 | 7,08 | 3,03 | FLOTILLIN3 (FLOT3) |
| AT1G18300 | 5,36 | 5,52 | 3,96 | 4,59 | 5,14 | 3,21 | 7,26 | 7,37 | 3,03 | NUDIX HYDROLASE HOMOLOG 4 (NUDT4) |
| AT2G27080 | 5,26 | 4,88 | 3,22 | 4,23 | 5,03 | 3,06 | 6,34 | 6,52 | 2,63 | NDR/HIN1-LIKE 13 (NHL13) |
| AT1G19020 | 5,21 | 6,52 | 6,32 | 5,89 | 6,84 | 3,50 | 6,84 | 7,87 | 3,90 | HYPOXIA RESPONSE UNKNOWN PROTEIN 35 (HUP35) |
| AT4G37290 | 5,20 | 5,32 | 8,32 | 5,61 | 6,42 | 4,10 | 6,42 | 8,14 | 5,35 | PRECURSOR OF PAMP-INDUCED PEPTIDE 2 (PREPIP2) |
| AT5G59820 | 5,20 | 6,55 | 5,86 | 4,13 | 4,78 | 3,29 | 6,03 | 7,39 | 4,45 | RESPONSIVE TO HIGH LIGHT 41 (RHL41; ZAT12) |
| AT1G07135 | 5,03 | 5,62 | 4,45 | 5,53 | 5,22 | 2,75 | 6,34 | 6,34 | 2,77 | glycine-rich protein |
| AT3G28340 | 4,95 | 5,43 | 4,47 | 5,03 | 5,52 | 3,06 | 6,24 | 6,44 | 3,25 | GALACTURONOSYLTRANSFERASE-LIKE 10 (GATL10) |
| AT2G32140 | 4,93 | 5,98 | 5,40 | 5,68 | 7,05 | 3,90 | 5,83 | 7,19 | 4,12 | transmembrane receptor |
| AT5G64310 | 4,87 | 5,83 | 4,28 | 4,62 | 6,73 | 3,43 | 5,77 | 6,33 | 3,28 | ARABINOGLACTAN PROTEIN 1 (AGP1) |
| AT5G41750 | 4,85 | 5,85 | 4,30 | 5,63 | 6,71 | 2,62 | 5,67 | 6,39 | 2,68 | Disease resistance protein (TIR-NBS-LRR class) family |
| AT2G32190 | 4,85 | 6,33 | 6,35 | 5,23 | 8,20 | 3,56 | 6,27 | 7,92 | 4,57 | CYSTEINE-RICH TRANSMEMBRANE MODULE 4 (ATHCYSTM4) |

**downregulated DEGs**
| ATG | WT<br>(log<sub>2</sub> FC) |  |  | *ahk2 ahk3*<br>(log<sub>2</sub> FC) |  |  | *cca1 lhy*<br>(log<sub>2</sub> FC) |  |  | short description |
| --- | --- | --- | --- | --- | --- | --- | --- | --- | --- | --- |
|  | 4 h | 6 h | 12 h | 4 h | 6 h | 12 h | 4 h | 6 h | 12 h |  |
| AT1G11130 | -1,03 | -1,56 | -2,78 | -1,11 | -1,94 | -3,65 | -1,64 | -2,48 | -6,30 | STRUBBELIG-RECEPTOR FAMILY 9 (SRF9);SCRAMBLED (SCM) |
| AT1G27360 | -1,12 | -1,48 | -1,87 | -1,15 | -1,82 | -2,11 | -1,48 | -2,37 | -2,31 | SQUAMOSA PROMOTER-LIKE 11 (SPL11) |
| AT4G07825 | -1,13 | -1,78 | -1,34 | -1,41 | -1,55 | -1,40 | -1,86 | -2,37 | -2,02 | transmembrane protein |
| AT3G18320 | -1,14 | -1,90 | -2,10 | -1,77 | -2,24 | -1,84 | -2,32 | -3,15 | -2,49 | F-box and associated interaction domains-containing protein |
| AT3G52170 | -1,28 | -1,63 | -1,69 | -1,12 | -1,36 | -1,66 | -1,30 | -2,08 | -2,69 | DNA binding protein |
| AT2G05995 | -1,47 | -2,04 | -3,93 | -2,12 | -1,69 | -3,71 | -2,24 | -2,65 | -4,38 | other_RNA |

To compare the transcriptional response of the different genotypes, the DEGs were compared for each time point (Figure 2). About two third of the responsive genes of WT were also found in the two more sensitive genotypes, which however had a higher total number of DEGs. Further, there is a significant and increasing number of DEGs characteristic for the stress-sensitive genotypes *ahk2 ahk3* and *cca1 lhy* (Supplemental Table S3) which also show a stronger regulation over time. One obvious difference to WR is the downregulation of numerous *SMALL AUXIN UP-RNA* (*SAUR*) genes. At the 6 h time point, 25 of the 40 known *SAUR* genes are strongly downregulated in *ahk2 ahk3* mutants and 6 *SAUR* genes in *cca1 lhy*. The number of downregulated *SAUR* genes and the degree of downregulation increases in both genotypes at the 12 h time point. Investigation of the functional relevance of these *SAUR* genes is an interesting direction for future research on the photoperiod stress syndrome.

**Figure 2.**
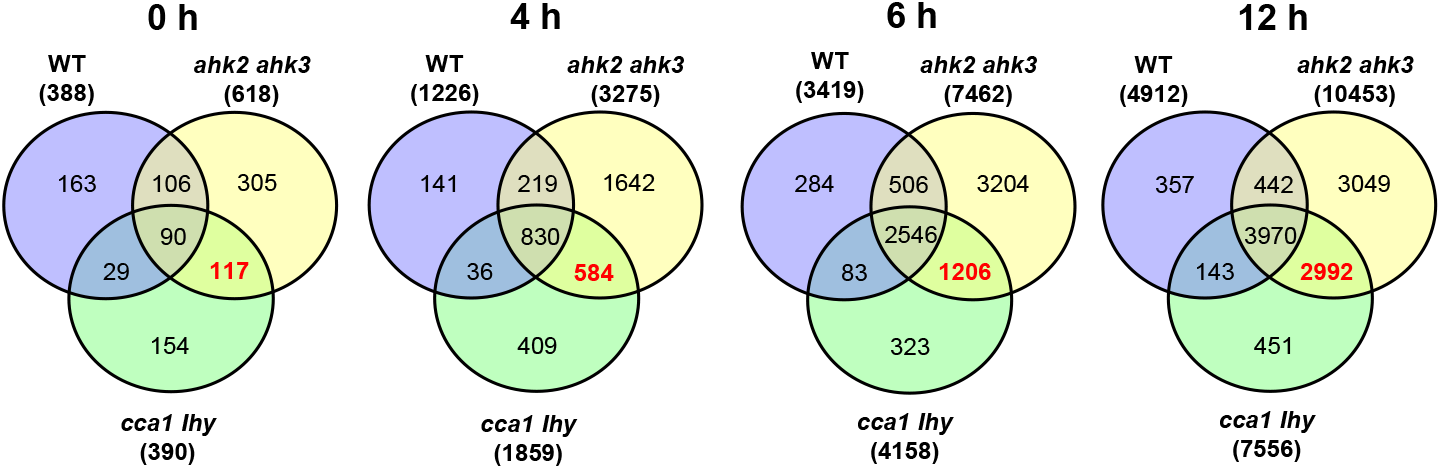
Overlap of DEGs of the different genotypes in response to photoperiod stress. Venn diagrams showing the overlap in DEGs between WT, *ahk2 ahk3* and *cca1 lhy* after photoperiod stress treatment at different time points. 5-weeks-old short-day grown plants were exposed to an extended light period of 32 h followed by a normal short-day night (see Fig. 1A). Numbers between brackets indicate the total number of DEGs (|fold-change| = 2; Bonferroni-corrected p-value ≤ 0.05) in PLP-treated plants compared to control plants at the respective time points. The bold red number in the Venn diagram indicates the number of DEGs occurring specifically in the stress-sensitive genotypes *ahk2 ahk3* and *cca1 lhy*.

### Numerous genes related to oxidative stress are responsive to photoperiod stress

As mentioned above, GO term analysis pointed to a strong correlation between photoperiod stress and oxidative stress (Figure 1E-G; Supplemental Table S1). Consistently, comparison of the top 20 strongest up- and down-regulated DEGs for the different genotypes (Supplemental Tables S4-S9) identified several commonly regulated genes related to oxidative stress among them *OXI1, RBOHC, PRX4, PRX37, ZAT11, CML37, CML38* and *ERF71*). The strong induction of these genes was confirmed for all three genotypes by qRT-PCR analysis (Figure 3, Supplemental Table S9). *OXI1* encodes a protein kinase necessary for oxidative burst-mediated signaling in *Arabidopsis* (Rentel et al., 2004). *RBOHC* is required for the production of ROS in response to an extracellular ATP stimulus (Kaya et al., 2019) and both *PRX4* and *PRX37* encode apoplastic oxidative burst peroxidases (Daudi et al., 2012; O’Brien et al., 2012; Valerio, De Meyer, Penel, & Dunand, 2004). The transcription factors (TFs) ZAT11 and ERF71 are involved in nickel ion tolerance (Liu et al., 2014) or hypoxia (Hess, Klode, Anders, & Sauter, 2011), respectively. Both transcription factor genes can be induced strongly by H_2_O_2_ (Hieno et al., 2019). The calmodulin-like proteins CML37 and CML38 are involved in drought stress and herbivory tolerance (S.S. Scholz, Reichelt, Vadassery, & Mithöfer, 2015; S. S. Scholz et al., 2014) or in hypoxia stress (Lokdarshi, Conner, McClintock, Li, & Roberts), respectively.

**Figure 3.**
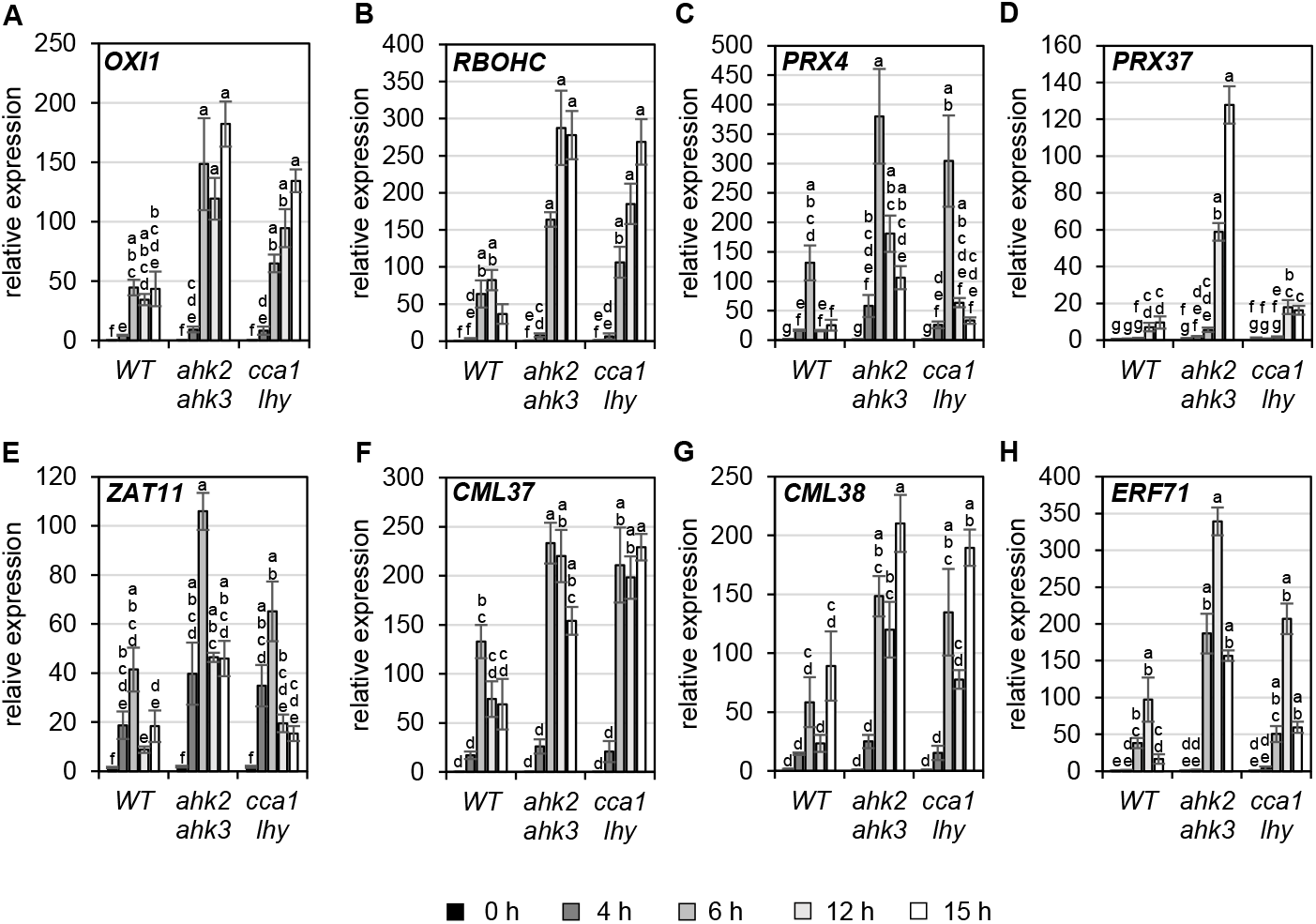
Genes related to oxidative stress are upregulated in all genotypes in response to photoperiod stress. Plants were grown under short day conditions for five weeks before exposure to a 32 h prolonged light period (PLP) (see Fig. 1A). (A - H) Transcript levels of *OXI1, RBOHC, PRX4, PRX37, ZAT11, CML37, CML28* and *ERF71* in WT, *ahk2 ahk3* and *cca1 lhy* plants at different time points during the night following PLP measured by qRT-PCR. Only results for the response to PLP-treatment are shown. An overview of all expression levels including control conditions is shown in Supplemental Table S10. All values are expressed relative to WT control at 0 h which was set to 1. Error bars represent SE (n ≥ 3). Letters indicate different statistical groups (p ≤ 0.05).

As both GO term analysis and DEGs in the top 20 revealed a strong connection to oxidative stress, the transcriptional regulation of genes encoding enzymes involved in the scavenging of ROS were analyzed in more detail. QT cluster analysis revealed three major clusters (Supplemental Figure S3, Supplemental Data 3). Cluster 1 shows an upregulation over time in all genotypes, which is much stronger in the stress-sensitive genotypes. *PRX34* belongs to this cluster which was previously identified as a photoperiod stress responsive gene involved in the oxidative burst (Abuelsoud et al., 2020). *CAT2* is one of the genes of cluster 2, which show a strong downregulation over time, which is even stronger in the stress-sensitive mutants. Cluster 3 consists of genes with a particularly strong expression at 4 h and 6 h time points for *ahk2 ahk3* mutants; *PRX4* belongs to this cluster.

These results indicate that photoperiod stress is associated with a strong regulation of oxidative stress-related genes. Oxidative stress is caused by an increase in reactive oxygen species be it H_2_O_2_, single oxygen or superoxide. Plants have evolved scavenging mechanisms to prevent cellular damage by these ROS generated under normal conditions or in stressful environments. Several core sets of ROS-responsive genes have been identified: Hieno et al. (2019) unraveled a core set of 60 H_2_O_2_-responsive TFs after H_2_O_2_ treatment; Zandalinas, Sengupta, Burks, Azad, and Mittler (2019) identified in response to short high light treatment 82 H_2_O_2_- and RBOHD-dependent genes and Lai et al. (2012) investigated the relation between the circadian clock and ROS-responsive genes and identified a core set of 167 genes of which 140 were clock-regulated. In addition, also transcriptional profiles specific for the response to H_2_O_2_, superoxide and singlet oxygen were identified (Gadjev et al., 2006). Specific transcriptional footprints should support the assessment of the functional roles of ROS in biological processes (Willems et al., 2016). The data sets of these studies were used to investigate the response profile of the oxidative stress-regulated genes during photoperiod stress.

The different core sets of ROS-responsive genes (Hieno et al. (2019), Zandalinas et al. (2019), Lai et al. (2012), Gadjev et al. (2006)) were first compared with each other (Supplemental Fig. S5A). Only a small overlap is observed between all four studies probably caused by the different experimental setups used to increase ROS production. Therefore, these sets of genes were pooled to form a new master core set of 283 ROS-responsive genes (Supplemental Data 4). QT clustering of transcript levels after photoperiod stress of this master core set of ROS-responsive genes indicated that there is a strong regulation of these genes starting at 4 h during the dark relaxation (Figure 4A). Four different clusters can be recognized: Cluster I with a strong time-dependent upregulation starting at the 4 h time point; cluster II with a strong time-dependent upregulation starting at the 6 h time point; cluster III showing a strong upregulation after 4 h and 6 h before a strong decrease in expression is visible at the 12 h time point; cluster IV with a time-dependent downregulation. Common for all clusters is the stronger response of the stress-sensitive mutants *ahk2 ahk3* and *cca1 lhy*. Transcript levels of representative genes of each cluster (*ZAT12, ERF1, CBF2* and *PRE1*)measured by qRT-PCR confirm the transcriptional regulation as shown in the heatmap (Figure 4B-E).

**Figure 4.**
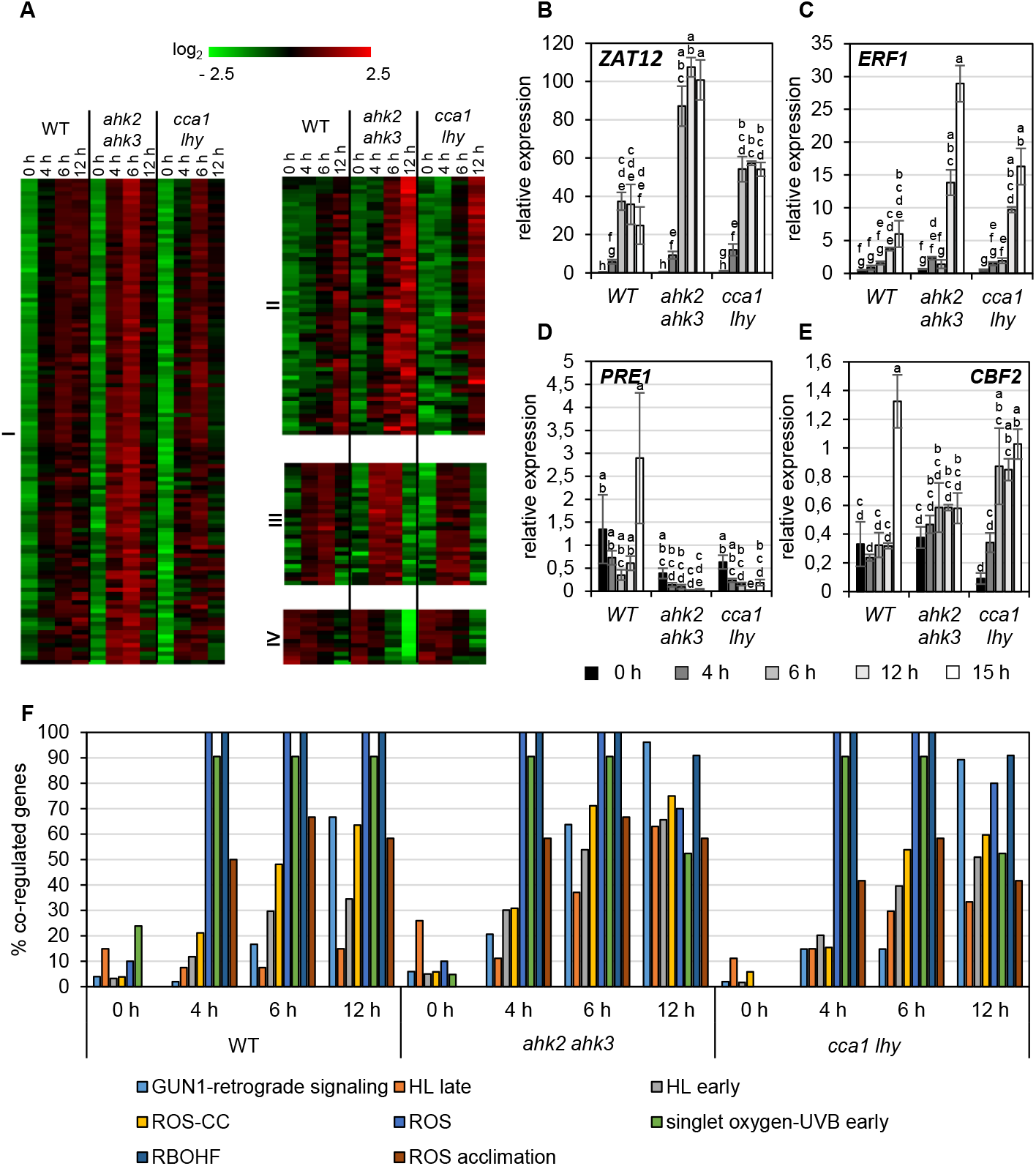
Photoperiod stress is associated with a strong transcriptional regulation of genes involved in oxidative stress. (A) QT clustering of genes encoding TFs identified by Hieno *et al*. (2019), Zandalinas *et al*. (2019), Lai *et al*. (2012) and Gadjev *et al*. (2006) as ROS-responsive genes. The overlap of the genes identified in the different studies is shown in Supplemental Figure S4. Four different clusters (I-IV) can be recognized. An overview of the regulation of these ROS regulated genes after PL (prolonged light) can be found in Supplemental Data 4. (B – E) Transcript levels of representative genes of each of the four cluster shown in A, i.e. *ZAT12* (B), *ERF1* (C), *PRE1* (D), and *CBF2* (E) in WT, *ahk2 ahk3* and *cca1 lhy* plants during the night following the 32 h PLP measured by qRT-PCR. Only results for PLP-treatment are shown. An overview of all expression levels including also control conditions can be found in Supplemental Table S10. All values are expressed relative to 0 h WT control which was set to 1. Error bars represent SE (n ≥ 3). Letters indicate different statistical groups. (F) Percentage of photoperiod stress-responsive genes that are co-regulated with the genes of the ROS wheel as defined by Willems *et al*. (2016). An overview of the regulation of these transcriptome profiles after photoperiod stress is given in Supplemental Data 6.

In addition, we analyzed the proportion of genes of the master core set regulated at different time points in the different genotypes after photoperiod stress (Supplemental Fig. S4C). Results clearly show that in all genotypes the proportion of regulated master core set ROS-responsive genes increase over time with the highest co-regulation at time points 6 h and 12 h. Furthermore, comparing the photoperiod stress-responsive transcriptome profile with the distinct superoxide-, singlet oxygen- and H_2_O_2_-induced gene profiles identified by Gadjev et al. (2006) (Supplemental Figure S4B) reveals a strong overlap with singlet oxygen-induced transcript profile (Supplemental Fig. S4B, Supplemental Data 5). Similarly, among the different ROS footprints identified by Willems et al. (2016) a strong overlap with singlet oxygen-UV-B early, RBOH-related and oxidative stress (ROS)-related transcript profiles (Figure 4F, Supplemental Data 6) can be recognized. Together, these results point to strong impact of ROS on the transcriptomic landscape in response to photoperiod stress.

### The transcriptional profile of photoperiod stress resembles transcriptional changes caused by pathogen and ozone stress

Photoperiod stress is a relative new form of abiotic stress and not much is known about similarities with other stresses. We therefore performed a meta-analysis comparing the transcriptomic profile caused by photoperiod stress with changes caused by other biotic and abiotic stresses including a shift to BL, drought stress, heat stress, cold stress, salt stress, ozone treatment, fluctuating light and high light stress (Ding et al., 2014; D. Huang, Wu, Abrams, & Cutler, 2008; J. Huang, Zhao, & Chory, 2019; Kleine, Kindgren, Benedict, Hendrickson, & Strand, 2007; Larkindale & Vierling, 2008; B. H. Lee, Henderson, & Zhu, 2005; Schneider et al., 2019; Tosti et al., 2006; Truman, Bennett, Kubigsteltig, Turnbull, & Grant, 2007) (Figure 5, Supplemental Data 8). In addition, we also compared our dataset with the transcriptome profile of circadian clock regulated genes (Covington et al., 2008) as a previous study found a connection to the circadian clock (Nitschke et al., 2016). At the 0 h time point, the number of co-regulated genes is relatively low in all genotypes which also reflects the fact that the photoperiod stress response starts later during the night following the PLP. This was also observed in the ROS-responsive transcript profile overlaps (Fig. 4; Supplemental Fig. S5). Most co-regulated genes can be found in studies with ozone treatment and pathogen attack in WT. Already after 4 h, approximately 37% of photoperiod stress regulated genes are identical to those regulated by ozone stress. The percentage of co-regulation even increases to almost 70% at time point 12 h. For pathogen attack, there is a co-regulation of more than 50% of the genes. In the stress-sensitive mutants, the co-regulation is also highest with ozone stress and pathogen attack. However, in these genotypes a higher overlap is found also with the transcriptional response to other stresses. The strong stress response of these mutants may probably activate general stress mechanisms which does not reveal specific information concerning the photoperiod stress response.

**Figure 5.**
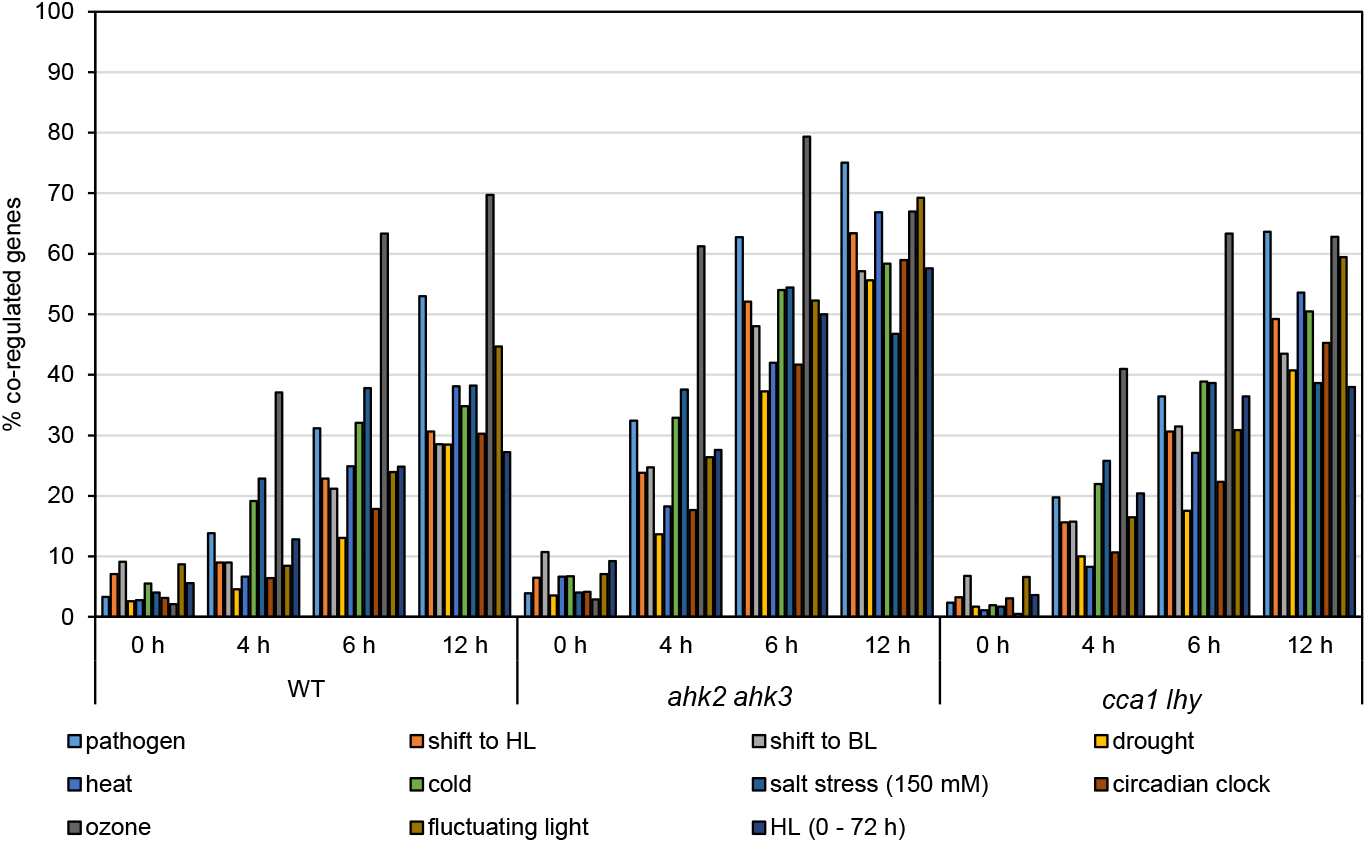
Photoperiod stress transcript profiles point to a similar response as in pathogen and ozone stress. Percent of responsive genes to photoperiod stress that are coregulated with genes responsive to various abiotic and biotic stresses. An overview of the photoperiod stress responses of the transcripts altered by these different stresses to photoperiod stress can be found in Supplemental Data 7.

### Photoperiod stress induces pathogen defense responses

The similarity of the transcriptional response to photoperiod stress and the response to pathogen attack (Figure 5) motivated us to explore links between these pathways in more detail. A central pathogen response pathway involves SA biosynthesis and signaling, which is required for SAR (Klessig, Choi, & Dempsey, 2018). Figure 6A shows hierarchical clustering of SA biosynthesis and signaling genes, which are listed in Supplemental Data 8. This analysis revealed that numerous SA-related genes are strongly deregulated during the night following a 32 h-prolongation of the light period. This change in the photoperiod causes a strong transcriptional upregulation especially at time point 6 h. Transcriptional regulation of selected SA biosynthesis and signaling genes has been confirmed by qRT-PCR for *EDS1, PAD4, PR1* and *FMO1* in WT and the stress-sensitive genotypes, *ahk2 ahk3* and *cca1 lhy* (Figure 6B-E).

**Figure 6.**
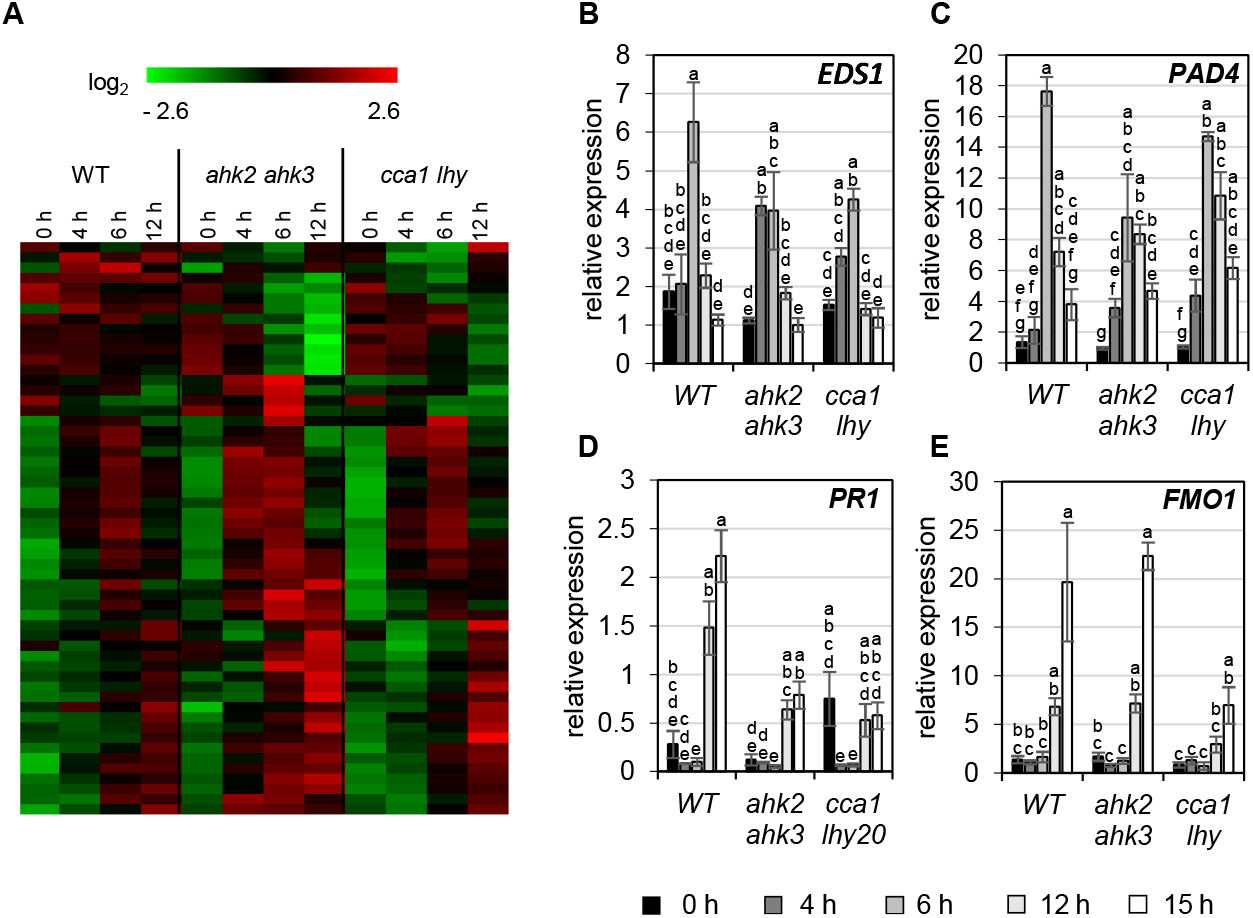
Photoperiod stress activates pathogen defense genes/responses. (A) Hierarchical clustering of genes involved in salicylic acid biosynthesis and signaling (Pearsons correlation). An overview of the regulation of these SA biosynthesis and signaling genes after photoperiod stress treatment is given in Supplemental Data 8. (B – E) Transcript levels of *EDS1* (B), *PAD4* (C), *SID2* (D), and *FMO1* (E) in WT, *ahk2 ahk3* and *cca1 lhy* plants during the night following the 32 h PLP measured by qRT-PCR. Only results for PLP treatment are shown. An overview of all expression levels including also control conditions can be found in Supplemental Table S10. All values are expressed relative to 0 h WT control which was set to 1. Error bars represent SE (n ≥ 2). Letters indicate different statistical groups (p ≤ 0.05).

To evaluate the functional relevance of the SA pathway for the photoperiod stress response, mutants of key steps in SA-dependent and -independent defense responses (*pad4, ics1, eds5, npr1-1* and *fmo1-1*) were exposed to PLP treatment and the photoperiod stress response was evaluated in terms of peroxide levels and photoperiod stress marker gene expression (Figure 7). Most mutants responded similar to WT. Also *sid2*, a mutant SA-deficient mutant responded similar as WT questioning the functional relevance of SA. However, *npr1-1* mutants showed a reduced responsiveness to photoperiod stress as they had low peroxide levels and strongly reduced marker gene expression compared to WT. These results indicate that NPR1 is required to sense and respond to photoperiod stress.

**Figure 7.**
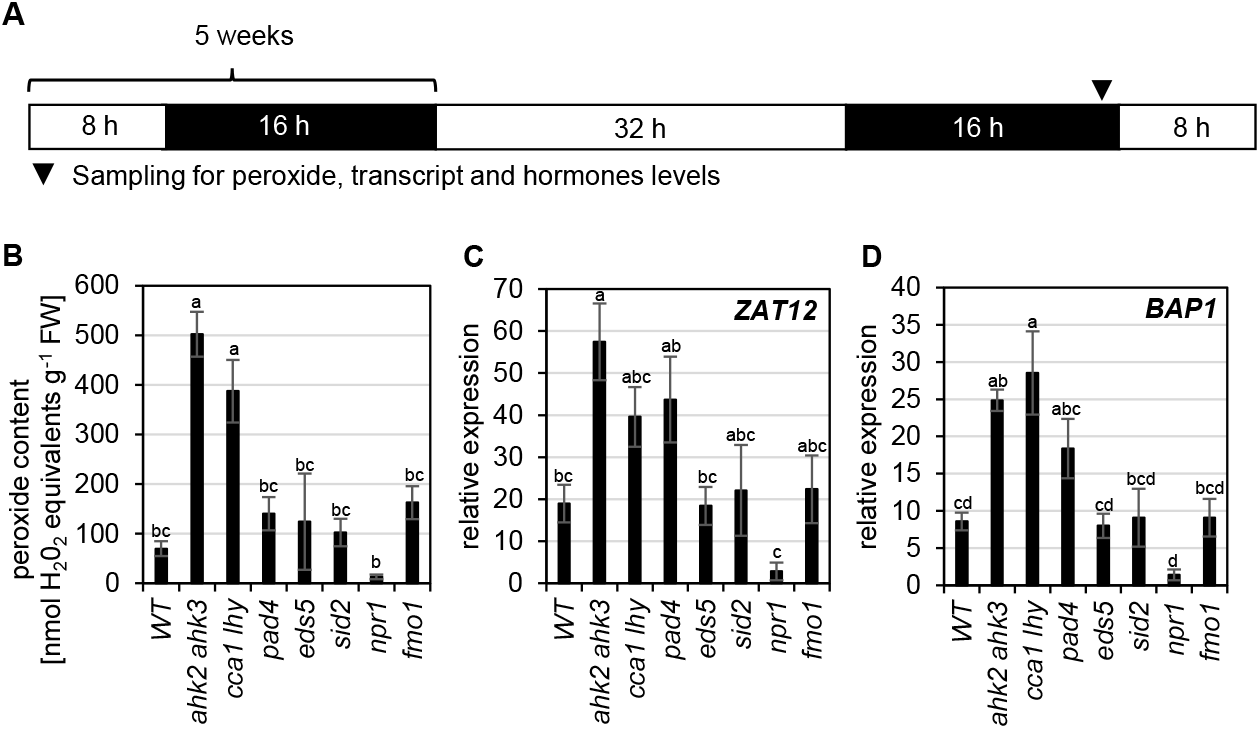
NPR1 has a central role during photoperiod stress. (A) Schematic overview of experimental setup. 5-weeks-old short-day grown plants were exposed to an extended light period of 32 h followed by a normal short-day night. Arrow head indicates sampling time point. (B) Peroxide levels after 32 h prolonged light period and (B-C) transcript levels of photoperiod stress marker genes *ZAT12* (C) and *BAP1* (D) in mutants of the SA-dependent and -independent defense response after pathogen infection. All values are expressed relative to 0 h WT control which was set to 1. Error bars represent SE (n ≥ 4). Letters indicate different statistical groups (p ≤ 0.05).

During pathogen defense responses, salicylic acid (SA) is an essential signaling molecule during both basal defense mechanisms and systemic acquired resistance (SAR). The formation of camalexin, which is one of the major phytoalexins in plant defense responses reducing the amount of bacteria in case an infection is present (Glawischnig, 2007) is induced by SA. Thus, SA and camalexin concentrations were measured at the end of the night in PLP-treated and control plants (WT only) (Figure 8A-B). After photoperiod stress, both SA and camalexin concentrations increase strongly.

**Figure 8.**
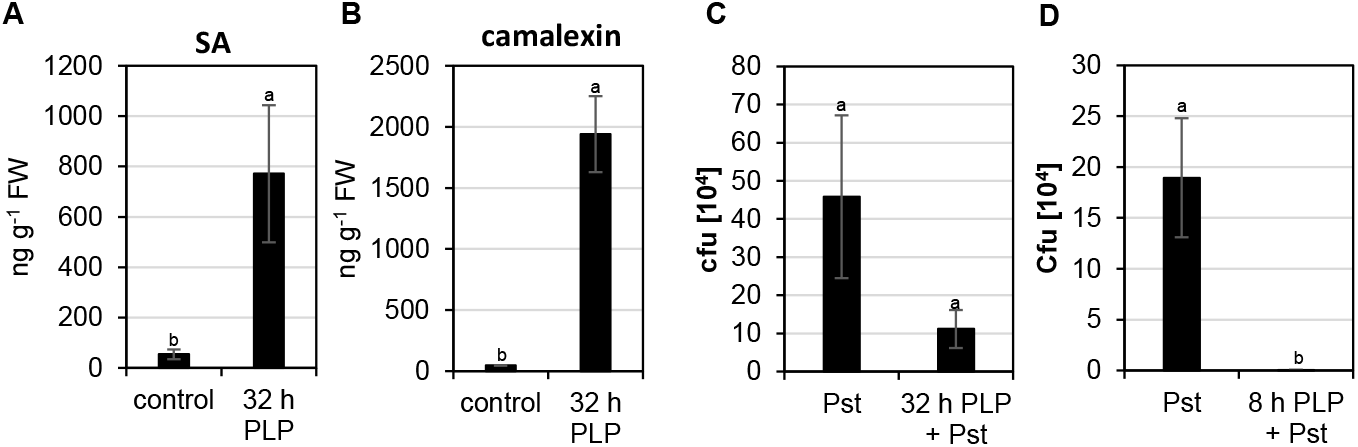
Photoperiod stress primes the plant pathogen response. 5-weeks-old short-day grown plants were exposed to an extended light period of 32 h followed by a normal short-day night. An overview of the experimental setup can be found in Figure 7A and arrowhead indicates sampling time point. (A) Salicylic acid (SA) and (B) camalexin concentration after photoperiod stress treatment. Error bars represent SE (n = 8). (C-D) Bacterial growth in *Arabidopsis* plants pretreated with a 32 h (C) or 8 h (D) prolonged light period. Bacterial infection was done during the day following the extended light period and bacteria were extracted from leaves 3 days later. Error bars represent SE (n = 8). Letters indicate different statistical groups (p≤ 0.05).

We hypothesized that because photoperiod stress activates similar responses as pathogen defense it might induce an increased plant immunity. To test this hypothesis, WT plants were inoculated with *Pseudomonas syringae* cv. *tomato* DC3000 (*Pst*) with and without previous exposure to photoperiod stress. We exposed WT plants to a photoperiod stress treatment of 32 h light before inoculation with *Pst*. Not yet fully developed leaves of control and photoperiod stress pretreated plants were inoculated with *Pst*, in the morning of the day after the photoperiod stress treatment. These leaves were chosen as they do not become flabby as mature leaves (Nitschke et al., 2016) which might interfere with the pathogen infection. In a separate experiment we reduced the duration of the photoperiod stress treatment to 8 h light prolongation thus avoiding the flabby phenotype in older leaves (7-10) before their inoculation with *Pst*. Colony forming units were counted three days post infection (dpi) and a reduced bacterial growth is observed in plants pretreated with photoperiod stress in both experimental scenarios (Figure 8C-D). These results suggest that photoperiod stress is able to induce an improved immunity in absence of pathogen.

## DISCUSSION

In this study, we have surveyed the transcriptomic changes occurring in response to strong photoperiod stress. A prolongation of the light period by 32 h resulted in massive transcriptomic changes during the night following the extended light period in WT *Arabidopsis* plants and even stronger alterations in the stress-sensitive *ahk2 ahk3* and *cca1 lhy* mutants (Fig. 1). Numerous transcriptional changes occur earlier than the development of the first visible photoperiod stress symptoms which start approximately 8 h after the prolongation of the light period has ended by the development of flabby leaves (Nitschke et al., 2016). Stronger changes in the transcriptional profile towards the end of the night coincide with the development of stronger photoperiod stress symptoms which eventually cause lesion formation (Nitschke et al., 2016; Nitschke et al., 2017).

Common to all genotypes was the prevalence of genes and GO terms related to oxidative stress among the the top 20 DEGs and top 5 GO terms (Figures 1 - 3, Table 1, Supplemental Tables S4-S9). Among them were a number of genes, e.g. *OXI1, RBOHC, PRX4* and *PRX37* that are known to be involved in the regulation of an oxidative burst after biotic or abiotic stresses (Daudi et al., 2012; Kaya et al., 2019; O’Brien et al., 2012; Rentel et al., 2004; Valerio et al., 2004). Similar regulation of the *PRX* genes was found by Abuelsoud et al. (2020). Also, the transcription factor genes *ZAT11* and *ERF71* are known to be strongly induced by H_2_O_2_ (Hieno et al., 2019). Moreover, photoperiod stress causes a strong transcriptional deregulation of genes coding for enzymes involved in the scavenging of reactive oxygen species (Supplemental Figure S4). These results suggested that the photoperiod stress display transcriptional responses similarities with transcriptomic changes caused by reactive oxygen species (ROS) which include H_2_O_2_, singlet oxygen and superoxide. This is consistent with the occurrence of oxidative stress in response to photoperiod stress as was shown by Abuelsoud et al. (2020). A core set of ROS-responsive genes was created based on comparisons of different studies investigating transcriptomic changes caused by ROS (Gadjev et al., 2006; Hieno et al., 2019; Lai et al., 2012; Zandalinas et al., 2019) (Supplemental Figure. S5, Supplemental Data 5, Figure. 4). Comparison with the photoperiod stress transcriptome revealed a strong regulation of these core ROS genes also after photoperiod stress. Additionally, the progression of the stress phenotype in the course of the night is reflected by the transcriptional profile as more ROS genes are deregulated towards the end of the night. Again, stronger responses were visible in the stress sensitive *ahk2 ahk3* and *cca1 lhy*. Comparison of the photoperiod stress transcriptome with transcriptional footprints for the assessment of the functional role of ROS during biological processes (Willems et al., 2016) also revealed a strong overlap with oxidative stress (ROS)-related transcript profiles (Figure 4; Supplemental Data 4). This is not unexpected as in a previous study (Abuelsoud et al., 2020), it was shown that one of the characteristics of photoperiod stress is the nightly increase in oxidative stress which is accompanied by the formation of peroxides. It was also revealed that the nightly ROS formation is associated with a strong increase of *PRX* gene induction and enhanced peroxidase activity and reduced catalase activity (Abuelsoud et al., 2020).

Meta-analysis comparing the photoperiod stress transcript profile with other biotic and abiotic stress response profiles revealed that there is a strong overlap with the effects of ozone and pathogen attack (Figure 4, Supplemental Data 8). These stresses also cause an apoplastic oxidative burst, similar as the nightly peak of peroxidases after photoperiod stress (Abuelsoud et al., 2020; Bolwell et al., 1999; Torres & Dangl, 2005; Van Breusegem & Dat, 2006). An apoplastic oxidative burst triggers salicylic acid (SA) biosynthesis and signaling. SA is a plant hormone with a well-established function during both basal (PTI) and induced (ETI) defense responses upon pathogen infection (for review: Herrera-Vasquez, Salinas, and Holuigue (2015)) and can act both as an antioxidant and as a potentiator of ROS. SA has also been shown to be involved in the regulation of responses to various abiotic stresses such as high light, heat, metal, drought, salinity, and cold stress (for review: Herrera-Vasquez et al. (2015); Khan, Fatma, Per, Anjum, and Khan (2015)).

During photoperiod stress genes involved in SA biosynthesis and signaling are strongly regulated resulting in SA-dependent and -independent defense responses (Figure 6, Supplemental Data 9). The strong induction of *EDS1* and *PAD4* in all genotypes, indicates that photoperiod stress induces a similar response as pathogen infection resulting in increased transcript levels of *SID2* which is also reflected in an increase in SA levels as shown in Nitschke et al. (2016) and in this study (Figure 7). In addition, camalexin levels are induced indicating that similar plant defense mechanisms are activated as during pathogen infection. Moreover, photoperiod stress pretreatment primes plants for a future pathogen infection as the amount of cfu dramatically reduced after photoperiod stress treatment (Fig. 7). This is a novel finding.

Among key components involved in defense responses after pathogen infection, NPR1 appears to be functional relevant for the photoperiod stress response (Figure 6) as mutants of these components showed an alleviation of the photoperiod stress. NPR1 is a master regulator of basal and systemic resistance in plants (Fu & Dong, 2013) and has been recently identified as part of a novel regulatory pathway in cold acclimation by interacting with HSFA1 factors (Olate, Jimenez-Gomez, Holuigue, & Salinas, 2018). Thus, photoperiod stress responses involve similar signaling components as are important during the SA-dependent defense responses (W. Huang, Wang, Li, & Zhang, 2020).

The fact that photoperiod stress results in a stronger SAR after pathogen infection in comparison to plants without previous photoperiod stress exposure (Figure 7) is a novel revolutionary fact about photoperiod stress, which might act as a protective mechanism in plant-defense responses. It is known that light is crucial for the activation of defense responses in plant-pathogen interactions (Ballaré, 2014; Delprato, Krapp, & Carrillo, 2015; Genoud, Buchala, Chua, & Metraux, 2002; Griebel & Zeier, 2008; Roberts & Paul, 2006; Roeber et al., 2020; Trotta, Rahikainen, Konert, Finazzi, & Kangasjarvi, 2014). Also the length of the light period affect the plant immune response as disease symptoms of *Arabidopsis* plants infected with Cauliflower Mosaic Virus are stronger in short day-grown plants than in plants grown under long day conditions (Cecchini et al., 2002). In addition, light quality influences the immune response (Ballaré, 2014; Fernandez-Milmanda et al., 2020). In view of this, we hypothesize that further characterization of the photoperiod stress response opens perspectives towards improving plant immunity by changing the light environment. Photoperiod stress might also be dependent on other light conditions such as the light quality and forms therefore an interesting starting point for future studies. The increasing use of LEDs (light emitting diodes) in greenhouses offers a possibility to use the light information optimally to increase growth-defense trade-offs in greenhouse cultures (Lazzarin et al., 2020) taking into account that photoperiod stress causes similar plant defense mechanisms as pathogen attack.

This study also identified some more candidate genes for future studies such as the *SAUR* genes which were also identified in a transcriptomic study investigating the response to high light stress (J. Huang et al., 2019). The *SAUR* regulation might be connected to the antagonistic function of auxin on the protectant cytokinin as it is known that an impairment of auxin perception in *ahk2 ahk3* reduces the stress sensitivity during photoperiod stress (Frank et al., 2021 - submitted). These findings classify the *SAUR* genes as good candidates for future functional studies.

## METHODS

### Plant Material and Growth Conditions

*Arabidopssi thaliana* accession Col-0 was used as WT. The following mutant and transgenic plants were used: *ahk2-5 ahk3-7* (Riefler, Novak, Strnad, & Schmulling, 2006), *cca1-1 lhy-20* (Nitschke et al., 2016), *npr1-1* (*Cao, Glazebrook, Clarke, Volko, & Dong, 1997*), *pad4* (Jirage et al., 1999)*, sid2/ics1* (Glazebrook, Rogers, & Ausubel, 1996), *fmo1-1* (Bartsch et al., 2006)*, eds5* (Nawrath, Heck, Parinthawong, & Metraux, 2002). Seeds were obtained from the European Arabidopsis Stock Centre (NASC). *Arabidopsis* plants were grown on soil for five weeks under short day (SD) conditions (8 h light/16 h darkness) in a growth chamber with light intensities of 100 to 150 μmol m^-2^ s^-1^, using a combination of Philips Son-T Agros 400W and Philips Master HPI-T Plus, 400W/645 lamps, at 22°C and 60% relative humidity.

### Stress Treatment

For stress treatments, five-weeks-old SD-grown plants were used. Photoperiod stress was induced by a 32 h light treatment (prolonged light period, PLP) followed by a normal 16 h night (Fig. 1A). Control plants remained under SD conditions. Stress parameters were analyzed in leaves 7-10. The harvest during the dark period was performed in green light.

### Plant Pathogen Infection

For the inoculation of *Arabidopsis* plants with *Pseudomonas syringae* cv. tomato DC3000 (*Pst*), the same method was used as described in Griebel and Zeier (2008).

### Analysis of Transcript Levels by RNA-seq and quantitative RT-PCR

Total RNA was extracted from leaf material (leaves 7-10). Only the distal parts of leaves 7-10 were harvested which is the most affected part of the leaves. Leaves were flash-frozen in liquid nitrogen and homogenized with a Retsch Mixer Mill MM2000 (Retsch, Haan, Germany). Total RNA was extracted using the NucleoSpin®RNA plant kit (Machery and Nagel, Düren, Germany) as described in the user’s manual or in Frank, Cortleven, Novak, and Schmulling (2020). For RNA-seq analysis, RNA was isolated from three biological samples at four different time points. The isolated RNA was send to BGI (Hongkong, China) and processed as described in (Cortleven, Ehret, Schmulling, & Johansson, 2019). In brief, a Nanodrop NA-1000 and a Bioanalyzer Agilent 2100 (Agilent Technologies, Santa Clara, CA, USA) were used to check RNA concentration, integrity and rRNA contamination. After DNase I treatment, mRNA was enriched by using oligo (dT) magnetic beads and fragmented into shorter fragments. First-strand cDNA was synthesized by using random hexamer primers, followed by second strand synthesis. After purification, end repair, and 3’ end single-nucleotide A (adenine) addition, sequence adaptors were ligated. Following PCR amplification and quality control by the Agilent 2100 Bioanalyzer and ABI StepOnePlus Real-Time PCR System (Thermo Fischer Scientific, Waltham, MA, USA), the library products were sequenced on the Illumina HiSeq™4000 platform. More than 22 million raw sequencing reads were obtained for each sample. After the removal of adaptors and low-quality reads, the obtained clean reads (approximately 21 million) were stored in FASTQ format (Cock, Fields, Goto, Heuer, & Rice, 2010). Sequences were aligned to the TAIR11 *Arabidopsis* reference genome using Bowtie2 (Langmead & Salzberg, 2012). Gene expression levels were quantified using RSEM (Li & Dewey, 2011) and differentially expressed genes (DEGs) were identified using the DESeq method (Love, Huber, & Anders, 2014) with the following default criteria: fold change ≥2 and Bonferroni correction (p-value ≤ 0.05). Gene Ontology (GO) annotation was performed using PANTHER (Mi, Muruganujan, Casagrande, & Thomas, 2013; Mi et al., 2019). Heatmaps were created using MEV (MultiExperiment Viewer (Howe, Sinha, Schlauch, & Quackenbush, 2011; Saeed et al., 2003)). For cluster analysis, MEV was used. Quality Threshold (QT) clustering was done using the following parameters: diameter: 0.7; minimum cluster size: 50; Euclidean distance or with a Pearsons correlation: minimum cluster size: 10, diameter: 0.3.

For quantitative real-time PCR (qRT-PCR), leaf material was collected at the same time points as for RNA-seq analysis. Quantitative real-time PCR analysis was performed as described in (Cortleven et al., 2019). Sequences of primers used for gene expression analysis are listed in Supplemental Table S11. Gene expression data were normalized against three or four different nuclear-encoded reference genes (*UBC21, TAFII15, PP2A* and/or *MCP2A*) according to (Vandesompele et al., 2002) and are expressed relative to the control treatment at time point 0 h which was set to 1.

### Determination of Peroxide Concentration

Water-soluble peroxides including hydrogen peroxide (H_2_O_2_) were determined as described in (Abuelsoud et al., 2020) by using a xylenol orange-based method (PierceTM Quantitative Peroxide Assay Kit (Aqueous), ThermoFischer Scientific, Berlin, Germany). Peroxides were extracted by the addition of 700 μl 0.1% ice cold trichloroacetic acid (TCA) to 100 mg finely ground and frozen leaf material. The 0.1% TCA inhibits catalase activity completely, while peroxidases are still active. After homogenization by vortexing, thawed homogenate was centrifuged at 10.000 *g* for 15 min at 4 °C. The concentration of peroxides was determined as described in the user’s manual. Hydrogen peroxide was used as standard and the watersoluble peroxide content was expressed as nmol H_2_O_2_ equivalent g^-1^ fresh weight (FW).

### Determination of SA and Camalexin Levels

SA and camalexin concentrations were determined as described in Valsamakis et al. (2020).

### Statistical Analysis

Statistical analyses were performed using SAS v.9.2 (SAS Institute GmbH, Heidelberg, Germany) or R studio (version 3.6.2). Data were analyzed by ANOVA followed by Tukey’s Post Hoc test. Normality and homogeneity of variance were tested using the Shapiro-Wilk and Levene tests (Neter *et al*., 1996). To meet the assumptions, datasets were transformed using log or square root transformations. When assumptions were not met, a non-parametric Kruskall-Wallis test was performed followed by a Wilcoxon-Mann-Whitney test to perform pairwise comparisons.

## SUPPLEMENTAL DATA

The following materials are available in the online version of this article.

**Supplemental Figure S1.** General overview of the RNA-seq dataset.

**Supplemental Figure S2.** QT clusters of significant genotype-dependent regulated genes.

**Supplemental Figure S3.** Photoperiod stress causes a strong transcriptional regulation of genes coding for enzymes involved in the scavenging of reactive oxygen species.

**Supplemental Figure S4.** Overlap between different datasets describing ROS-responsive transcription factors.

**Supplemental Table S1.** GO term enrichment in photoperiod stress-treated versus control plants.

**Supplemental Table S2.** Core-set of photoperiod stress-responsive genes.

**Supplemental Table S3.** Changes of transcript levels (log^2^ fold change) specific for the stress sensitive genotypes *ahk2 ahk3* and *cca1 lhy*.

**Supplemental Table S4.** Top 20 upregulated genes at different time points after photoperiod stress treatment in WT in comparison to untreated control.

**Supplemental Table S5.** Top 20 downregulated genes at different time points after photoperiod stress treatment in WT in comparison to untreated control.

**Supplemental Table S6.** Top 20 upregulated genes at different time points after photoperiod stress treatment in *ahk2 ahk3* in comparison to untreated *ahk2 ahk3*.

**Supplemental Table S7.** Top 20 downregulated genes at different time points after photoperiod stress treatment in *ahk2 ahk3* in comparison to untreated *ahk2 ahk3*.

**Supplemental Table S8.** Top 20 upregulated genes at different time points after photoperiod stress treatment in *cca1 lhy* in comparison to untreated *cca1 lhy*.

**Supplemental Table S9.** Top 20 downregulated genes at different time points after photoperiod stress treatment in *cca1 lhy* in comparison to untreated *cca1 lhy*.

**Supplemental Table S10.** Transcript levels of selected genes as determined by qRT-PCR normalized to WT control at 0 h.

**Supplemental Table S11.** Sequences of primers used for qRT-PCR.

**Supplemental Data 1.** Gene identifiers (ATGs) of statistically significant DEGs at different time points following photoperiod stress treatment in WT, *ahk2 ahk3* and *cca1 lhy*.

**Supplemental Data 2.** Expression of *Arabidopsis* genes identified in this RNA-seq analysis at different time points after photoperiod stress treatment in WT, *ahk2 ahk3* and *cca1 lhy* expressed relative to control treatment.

**Supplemental Data 3.** Transcript profiles of genes encoding ROS scavenging enzymes after photoperiod stress.

**Supplemental Data 4.** Changes is transcript abundance of ROS-responsive genes after photoperiod stress.

**Supplemental Data 5.** Overlap of the response to photoperiod stress with H_2_O_2_-, superoxide- and singlet oxygen-specific transcriptome profiles.

**Supplemental Data 6.** Genes of the ROS wheel (Willems *et al*. (2016)) that respond to photoperiod stress.

**Supplemental Data 7.** The response to photoperiod stress of genes induced after different abiotic and biotic stress treatments.

**Supplemental Data 8.** Transcript profile of SA biosynthesis and signaling genes after photoperiod stress.

## ACKNOWLEDGEMENTS

We are grateful to Heidrun Haeweker for excellent technical assistance and Thomas Griebel for providing the *Pseudomonas syringae* strain. We acknowledge funding by Deutsche Forschungsgemeinschaft to T.S. (Schm 814/27-1 and Sfb 973).

---

**Supplemental Figure S1. General overview of the RNA-seq dataset.**

(A) PCA of the biological samples. Control samples cluster together (red circle). A strong diversification of light-treated samples is visible with a strong separation of the different genotypes at time point 12 h (blue circles). (B-C) QT clustering of differentially expressed genotype-regulated genes. 52% of all significantly regulated genotype-dependent genes belong to cluster 1 (B) and cluster 2 (C). An overview of the other clusters can be found in Supplemental Figure S2. (D-E) GO term analysis of genes belonging to cluster 1 (D) and to cluster 2 (E).

**Supplemental Figure S2. QT clusters of significant genotype-dependent regulated genes.**

QT clustering of differentially expressed genotype-regulated genes.

**Supplemental Figure S3. Photoperiod stress causes a strong transcriptional regulation of genes coding for enzymes involved in the scavenging of reactive oxygen species.**

5-weeks-old short day-grown plants were exposed to an extended light period of 32 h followed by a normal short-day night. The experimental setup is shown in Fig. 1A. (A) QT clustering of genes encoding enzymes involved in the scavenging of reactive oxygen species (Pearsons correlation). Three clusters are found: cluster 1 showing an upregulation over time, cluster 2 showing a downregulation over time and cluster 3 showing a strong expression at time points 4 h and 6 h. Depicted are the ratios of PLP versus control plants for each genotype and time point. A complete list of the genes coding for scavenging enzymes used in this analysis including their expression levels is shown in Supplemental Data 3.

**Supplemental Figure S4. Overlap of photoperiod stress responsive transcription factor genes with different datasets describing ROS-responsive transcription factor genes.**

(A) Venn diagram showing the overlap between the ROS-responsive genes identified by Hieno *et al*. (2019), Zandalinas *et al*. (2019), Lai *et al*. (2012) and Gadjev *et al*. (2006). (B) Venn diagrams showing the proportion of the 283 ROS-responsive genes shown in (A) that are induced or repressed by photoperiod stress in the different genotypes at different time points. (C) Overlap of H_2_O_2_, superoxide and singlet oxygen specific transcript profiles as identified by Gadjev *et al*. (2006). (D) Percentage of DEGs induced by photoperiod stress co-regulated with genes responsive to H_2_O_2_, superoxide and singlet oxygen shown in (C). An overview of the regulation of these transcripts after photoperiod stress is given in Supplemental Data 4 and in Supplemental Data 5.

