## Supplemental Table S4-S9 for "The transcriptomic landscape of the photoperiodic stress response in *Arabidopsis thaliana* resembles the response to pathogen infection"

**Supplemental Table S4. Top 20 upregulated genes at different time points after photoperiod stress treatment in WT in comparison to untreated control.** FC, fold change; p-value according to Bonferroni.

| **Time point 0 h** | |  |  |
| --- | --- | --- | --- |
| ATG | log_2_ FC | p-value | short description |
| AT3G01270 | 16.0116088 | 0.00011323 | pectate lyase family protein |
| AT5G42800 | 7.99419411 | 1.63E-08 | dihydroflavonol reductase (DFR) |
| AT5G13930 | 7.57010341 | 1.85E-27 | encodes chalcone synthase (CHS), a key enzyme involved in the biosynthesis of flavonoids (TT4) |
| AT4G22880 | 6.74289622 | 2.41E-07 | encodes leucoanthocyanidin dioxygenase (LDOX) (ANS) (TT18) (TDS4) |
| AT1G65480 | 6.67731583 | 1.63E-05 | FT |
| AT3G29590 | 6.64369572 | 3.89E-05 | encodes a malonyl-CoA:anthocyanidin 5-O-glucoside-6"-O-malonyltransferase (At4g14090) (AT5MAT) |
| AT5G05270 | 6.12327217 | 7.91E-31 | chalcone-flavanone isomerase family protein CHIL |
| AT1G01060 | 6.02381567 | 3.60E-44 | LHY encodes a myb-related putative transcription factor involved in circadian rhythm (LHY) |
| AT4G34410 | 5.92898149 | 0.00019282 | encodes a member of the ERF (ethylene response factor) subfamily B-3 of ERF/AP2 transcription factor family (RRTF1) |
| AT5G54060 | 5.64577102 | 6.66E-05 | encodes a anthocyanin 3-O-glucoside: 2"-O-xylosyl-transferase (UF3GT) |
| AT1G34580 | 5.23719617 | 0.00334224 | major facilitator superfamily protein |
| AT3G51240 | 5.23384093 | 1.37E-18 | encodes flavanone 3-hydroxylase that is coordinately expressed with chalcone synthase and chalcone isomerases (F3H) |
| AT5G07990 | 5.22763424 | 2.58E-22 | required for flavonoid 3' hydroxylase activity (TT7) |
| AT1G70260 | 5.16922649 | 1.20E-11 | encodes an endoplasmic reticulum (ER)-localized nodulin MtN21-like transporter family protein that negatively regulates resistance against biotrophic pathogens (UMAMIT36) |
| AT2G44840 | 4.89521633 | 9.90E-05 | encodes a member of the ERF (ethylene response factor) subfamily B-3 of ERF/AP2 transcription factor family (ERF13) |
| AT3G22840 | 4.79486754 | 1.78E-19 | encodes an early light-inducible protein (ELIP1) |
| AT1G65060 | 4.65894167 | 2.62E-18 | encodes an isoform of 4-coumarate:CoA ligase (4CL) (4CL3) |
| AT4G09820 | 4.64596061 | 0.00012505 | TT8 is a regulation factor of flavonoid pathways (TT8) |
| AT5G17220 | 4.55373574 | 0.00909057 | encodes glutathione transferase belonging to the phi class of GSTs |
| AT1G76650 | 4.31819303 | 0.00536724 | CALMODULIN-LIKE 38 (CML38) |
| **Time point 4 h** | |  |  |
| ATG | log_2_ FC | p-value | short description |
| AT2G47040 | 35.5475897 | 2.99E-28 | share high homologies with a group of pectin methylesterases (PME), pollen specific, and is required for enhancing the growth of pollen tube in style and transmitting tract tissues (VGD1) |
| AT3G01270 | 17.243004 | 1.37E-06 | pectate lyase family protein |
| AT3G07820 | 15.7379915 | 0.03363671 | pectin lyase-like superfamily protein |
| AT1G56250 | 8.59114713 | 7.27E-14 | encodes an F-box protein that can functionally replace VirF, regulating levels of the VirE2 and VIP1 proteins via a VBF-containing SCF complex (VBF) |
| AT1G14540 | 8.51511118 | 7.48E-13 | peroxidase superfamily protein (PRX4) |
| AT4G31950 | 8.23091702 | 0.02038203 | member of CYP82C (CYP82C3) |
| AT4G30280 | 8.17045477 | 4.77E-15 | encodes a xyloglucan endotransglucosylase/hydrolase (ATXTH18) |
| AT1G01060 | 7.69842077 | 6.39E-59 | LHY encodes a myb-related putative transcription factor involved in circadian rhythm (LHY1) |
| AT3G02840 | 7.62217611 | 3.22E-17 | ARM repeat superfamily protein |
| AT1G65480 | 7.61906864 | 1.62E-05 | FT |
| AT3G46080 | 7.41646739 | 2.01E-10 | C2H2-type zinc finger family protein |
| AT3G25250 | 7.06062165 | 9.34E-11 | Arabidopsis protein kinase The mRNA is cell-to-cell mobile (OXI1) |
| AT5G42380 | 6.91406443 | 3.31E-13 | calmodulin like 37 (CML37) |
| AT3G22840 | 6.86397046 | 6.34E-06 | encodes an early light-inducible protein (ELIP1) |
| AT5G60350 | 6.85021253 | 0.01866566 | hypothetical protein |
| AT3G46090 | 6.80471126 | 9.24E-13 | C2H2 and C2HC zinc fingers superfamily protein (ZAT7) |
| AT2G02340 | 6.78023257 | 0.032307 | phloem protein 2-B8 (PP2-B8) |
| AT2G46830 | 6.72283626 | 3.59E-61 | CCA1 |
| AT2G37430 | 6.6743247 | 3.56E-11 | encodes a member of the zinc finger family of transcriptional regulators (ZAT11) |
| AT1G76650 | 6.67373567 | 4.96E-12 | calmodulin-like 38 (CML38) |

**Supplementary Table S4 (*continued*). Top 20 upregulated genes at different time points after photoperiod stress treatment in WT in comparison to untreated control.**

| **Time point 6 h** | |  |  |
| --- | --- | --- | --- |
| ATG | log_2_ FC | p-value | short description |
| AT2G47040 | 36.7866099 | 1.88E-32 | share high homologies with a group of pectin methylesterases (PME), pollen specific, and is required for enhancing the growth of pollen tube in style and transmitting tract tissues (VGD1) |
| AT2G47520 | 10.7947558 | 1.34E-07 | encodes a member of the ERF (ethylene response factor) subfamily B-2 of ERF/AP2 transcription factor family (ERF71) |
| AT5G66890 | 10.3445662 | 7.90E-09 | RPW8 -CNL gene (NRG1.3) |
| AT5G37490 | 9.34349215 | 1.15E-08 | RM repeat superfamily protein |
| AT4G30280 | 9.33950922 | 9.38E-21 | encodes a xyloglucan endotransglucosylase/hydrolase (XTH18) |
| AT5G51060 | 9.32103368 | 8.67E-15 | RHD2 (along with RHD3 and RHD4) is required for normal root hair elongation (RBOHC) |
| AT1G56240 | 9.02989994 | 1.90E-07 | phloem protein 2-B13 (PP2-B13) |
| AT4G28460 | 8.9948649 | 8.22E-11 | transmembrane protein |
| AT1G13480 | 8.97591712 | 7.36E-10 | hypothetical protein (DUF1262) |
| AT5G56960 | 8.8869028 | 2.78E-23 | basic helix-loop-helix (bHLH) DNA-binding family protein |
| AT2G21910 | 8.77614396 | 1.73E-05 | member of CYP96A (CYP96A5) |
| AT5G27765 | 8.66119758 | 8.69E-06 | transmembrane protein |
| AT1G30370 | 8.6558486 | 1.45E-08 | encodes a mitochondria-localized class III phospholipase A1 that plays a role in seed viability (DLAH) |
| AT4G08555 | 8.4865357 | 4.02E-21 | hypothetical protein;(source:Araport11) |
| AT2G37430 | 8.45685964 | 1.73E-14 | encodes a member of the zinc finger family of transcriptional regulators (ZAT11) |
| AT3G02840 | 8.4293975 | 8.13E-22 | ARM repeat superfamily protein |
| AT3G25250 | 8.33040095 | 4.28E-16 | Arabidopsis protein kinase The mRNA is cell-to-cell mobile (OXI1) |
| AT1G73550 | 8.30814101 | 5.59E-05 | encodes a Protease inhibitor/seed storage/LTP family protein |
| AT5G42380 | 8.25861505 | 5.75E-20 | calmodulin like 37 (CML37) |
| AT1G14540 | 8.19503335 | 1.40E-12 | peroxidase superfamily protein (PRX4) |
| **Time point 12 h** | |  |  |
| ATG | log_2_ FC | p-value | short description |
| AT1G66550 | 23.0516942 | 3.97E-22 | member of WRKY Transcription Factor; Group III (WRKY67) |
| AT5G55150 | 11.0801732 | 4.53E-08 | F-box SKIP23-like protein (DUF295) (ATFDR2) |
| AT2G23270 | 10.3180304 | 6.24E-09 | transmembrane protein |
| AT5G40990 | 10.3043815 | 2.22E-08 | component of plant resistance (GLIP1) |
| AT2G04040 | 10.1515971 | 8.52E-17 | AtDTX1 (At2g04040) has been identified as a detoxifying efflux carrier for plant-derived antibiotics and other toxic compounds, including Cd2+ (DTX1) |
| AT1G17180 | 9.89435514 | 9.91E-07 | encodes glutathione transferase belonging to the tau class of GSTs.(GSTU25) |
| AT1G05680 | 9.69999127 | 1.02E-09 | encodes a UDP-glucosyltransferase (UGT74E2) |
| AT2G47520 | 9.62907877 | 1.18E-08 | encodes a member of the ERF (ethylene response factor) subfamily B-2 of ERF/AP2 transcription factor family (ERF71) |
| AT3G54530 | 9.52525977 | 0.04614494 | hypothetical protein |
| AT5G57123 | 9.50438982 | 7.32E-10 | hypothetical protein |
| AT3G25490 | 9.42236823 | 0.00037453 | protein kinase family protein |
| AT2G41230 | 9.35139775 | 6.61E-09 | encodes an ER-localized plant hormone-responsive gene and appears to act redundantly with ARGOS and ARL during organ growth (OSR1) |
| AT1G53990 | 9.13856848 | 7.14E-08 | contains lipase signature motif and GDSL domain (GLIP3) |
| AT3G44830 | 9.06894604 | 5.95E-09 | lecithin:cholesterol acyltransferase family protein |
| AT4G37390 | 9.02223692 | 3.79E-14 | encodes an IAA-amido synthase that conjugates Asp and other amino acids to auxin in vitro (BRU6) |
| AT2G44070 | 9.01163022 | 9.54E-07 | NagB/RpiA/CoA transferase-like superfamily protein |
| AT5G36970 | 9.00968942 | 3.03E-05 | NDR1/HIN1-like protein, expression induced during incompatible response to a pathogen, expression is at least partly dependent on the salicylic acid signaling pathway (NHL25) |
| AT2G21910 | 8.96363027 | 7.50E-06 | member of CYP96A (CYP96A5) |
| AT4G08770 | 8.91254705 | 9.77E-07 | encodes a putative apoplastic peroxidase Prx37 (Prx37) |
| AT4G28460 | 8.90010554 | 1.36E-12 | transmembrane protein |

**Supplemental S5. Top 20 downregulated genes at different time points after photoperiod stress treatment in WT in comparison to untreated control.** FC, fold change; p-value according to Bonferroni.

| **Time point 0 h** | |  |  |
| --- | --- | --- | --- |
| ATG | log_2_ FC | p-value | short description |
| AT1G66550 | -18.9209283 | 5.16E-12 | member of WRKY Transcription Factor (WRKY67) |
| AT2G05540 | -6.03050553 | 1.61E-11 | glycine-rich protein family |
| AT5G61160 | -5.73567161 | 6.02E-06 | anthocyanin 5-aromatic acyltransferase 1 (AACT1) |
| AT4G16260 | -4.95055392 | 0.00476861 | encodes a putative beta-1,3-endoglucanase that interacts with the 30C02 cyst nematode effector. |
| AT5G57550 | -4.9229028 | 2.34E-09 | xyloglucan endotransglycosylase-related protein (XTR3) |
| AT4G36570 | -3.31414159 | 6.81E-11 | RAD-like 3 (RL3) |
| AT3G51400 | -3.24830689 | 8.68E-15 | hypothetical protein (DUF241) |
| AT2G33830 | -3.24364612 | 2.93E-11 | Dormancy/auxin associated family protein (DRM2) |
| AT1G15125 | -2.94075755 | 0.00514041 | S-adenosyl-L-methionine-dependent methyltransferases superfamily |
| AT4G06746 | -2.63914249 | 2.15E-06 | encodes a member of the DREB subfamily A-5 of ERF/AP2 transcription factor family (RAP2.9) |
| AT3G01960 | -2.54023285 | 1.66E-08 | hypothetical protein |
| AT1G02460 | -2.51829554 | 0.0067568 | pectin lyase-like superfamily protein |
| AT2G36270 | -2.47301058 | 0.0003883 | encodes a member of the basic leucine zipper transcription factor family (ABI5) |
| AT5G40180 | -2.34403758 | 0.020447 | Pmr5/Cas1p GDSL/SGNH-like acyl-esterase family protein |
| AT2G46310 | -2.31977785 | 7.09E-10 | CRF5 encodes one of the six cytokinin response factors. It is transcriptionally upregulated in response to cytokinin (CRF5) |
| AT2G47870 | -2.26824555 | 0.00030782 | encodes a member of the CC-type glutaredoxin (ROXY) family that has been shown to interact with the transcription factor TGA2 and suppress ORA59 promoter activity (ROXY5) |
| AT3G32980 | -2.26684453 | 0.00010194 | unknown |
| AT3G05660 | -2.24278836 | 1.43E-07 | receptor like protein 33 (RLP33) |
| AT5G62360 | -2.20065674 | 2.98E-06 | pectin methylesterase inhibitor expressed throughout the plant (PMEI13) |
| AT5G23660 | -2.15357548 | 3.46E-09 | encodes a member of the SWEET sucrose efflux transporter family proteins (SWEET12) |
| **Time point 4 h** | |  |  |
| ATG | log_2_ FC | p-value | short description |
| AT1G66550 | -19.4193265 | 2.66E-13 | member of WRKY Transcription Factor (WRKY67) |
| AT3G22231 | -3.13446446 | 0.01156885 | encodes a member of a novel 6 member Arabidopsis gene family (PCC1) |
| AT2G43590 | -2.40847766 | 9.04E-05 | chitinase family protein |
| AT5G59662 | -2.28490665 | 3.71E-07 | natural antisense transcript overlaps with AT5G59660 |
| AT4G25630 | -2.09608716 | 1.53E-06 | encodes a fibrillarin, a key nucleolar protein in eukaryotes which associates with box C/D small nucleolar RNAs (snoRNAs) directing 2'-O-ribose methylation of the rRNA (FIB2) |
| AT4G26150 | -2.05259461 | 9.75E-05 | GATA TRANSCRIPTION FACTOR 22 (GATA22) |
| AT3G23830 | -2.01453448 | 9.09E-11 | GLYCINE-RICH RNA-BINDING PROTEIN 4 (GRP4) |
| AT1G32583 | -2.01434853 | 0.00135447 | microRNA (MIR400) is derived from the first intron of At1g32583 in the 5'UTR |
| AT5G27120 | -1.98609504 | 1.16E-14 | SAR DNA-binding protein, putative, strong similarity to SAR DNA-binding protein-1 (Pisum sativum) GI:3132696; |
| AT2G38400 | -1.85524229 | 0.00032398 | GLYOXYLATE AMINOTRANSFERASE 3 (AGT3) |
| AT3G59670 | -1.85512064 | 1.93E-08 | elongation factor |
| AT1G67120 | -1.85333321 | 1.75E-15 | represents a homolog of the yeast MDN gene, which encodes a non-ribosomal protein involved in the maturation and assembly of the 60S ribosomal subunit; MIDASIN 1 (MDN1) |
| AT1G78930 | -1.84438189 | 2.09E-06 | mitochondrial transcription termination factor family protein |
| AT3G61920 | -1.84432363 | 0.04047352 | UvrABC system protein C |
| AT4G12830 | -1.84310817 | 4.92E-05 | alpha/beta-Hydrolases superfamily protein |
| AT2G47880 | -1.83059932 | 0.00010139 | encodes a member of the CC-type glutaredoxin (ROXY) family. CEP DOWNSTREAM 2 (CEPD2; ROXY9) |
| AT5G14580 | -1.82270387 | 3.84E-10 | polyribonucleotide nucleotidyltransferase |
| AT1G30960 | -1.8059281 | 0.00070549 | ortholog of ERA (E. coli RAS-like protein)-related GTPase (ERG); E. COLI RAS-LIKE PROTEIN)-RELATED GTPASE 2 (ERG2) |
| AT3G44750 | -1.80546554 | 3.60E-09 | encodes a histone deacetylase; HISTONE DEACETYLASE 3 (HDA3) |
| AT4G36410 | -1.80276736 | 0.0052041 | ubiquitin-conjugating enzyme; UBIQUITIN-CONJUGATING ENZYME 17 |

**Supplementary Table S5 (*continued*). Top 20 downregulated genes at different time points after photoperiod stress treatment in WT in comparison to untreated control.**

| **Time point 6 h** | |  |  |
| --- | --- | --- | --- |
| ATG | log_2_ FC | p-value | short description |
| AT1G01590 | -3.81879508 | 4.45E-06 | FERRIC REDUCTION OXIDASE 1 (FRO1) |
| AT4G26530 | -3.67419898 | 3.19E-15 | FRUCTOSE-BISPHOSPHATE ALDOLASE 5 (FBA5) |
| AT5G59662 | -3.33044717 | 4.44E-08 | natural antisense transcript overlaps with AT5G59660; antisense_long_noncoding_rna |
| AT5G04150 | -3.16312321 | 0.00052934 | encodes a member of the basic helix-loop-helix transcription factor family protein (BHLH101) |
| AT5G51310 | -3.08025496 | 2.83E-06 | mutants exhibit longer root hairs under phosphate-deficient conditions |
| AT3G01960 | -3.07481756 | 1.22E-11 | hypothetical protein |
| AT5G23660 | -2.92122976 | 4.19E-19 | encodes a member of the SWEET sucrose efflux transporter family proteins (SWEET12) |
| AT4G12830 | -2.86063433 | 8.77E-16 | alpha/beta-Hydrolases superfamily protein |
| AT3G51400 | -2.80585279 | 0.00142214 | hypothetical protein (DUF241) |
| AT4G12320 | -2.77010958 | 3.61E-09 | member of CYP706A (CYP706A6) |
| AT1G32583 | -2.76515359 | 1.95E-07 | a microRNA MIR400 is derived from the first intron of At1g32583 in the 5'UTR |
| AT4G25780 | -2.71832438 | 9.22E-09 | CAP (Cysteine-rich secretory proteins, Antigen 5, and Pathogenesis-related 1 protein) superfamily protein (ATCAPE2) |
| AT1G78930 | -2.71502129 | 5.41E-13 | mitochondrial transcription termination factor family protein |
| AT1G07050 | -2.70973069 | 1.74E-26 | FITNESS encodes a protein with a single CCT domain and belongs to the CCT motif family genes (CMF); acts upstream JUB1 thereby controlling H2O2 levels (FITNESS) |
| AT5G05890 | -2.67020445 | 7.83E-23 | encodes a nicotinate-N-glycosyltransferase, UDP-GLUCOSYL TRANSFERASE 76C5 (UGT76C5) |
| AT1G52342 | -2.63814252 | 5.33E-10 | hypothetical protein |
| AT5G58370 | -2.63460446 | 1.18E-35 | P-loop containing nucleoside triphosphate hydrolases superfamily protein; ENGB-3 |
| AT1G56710 | -2.61978371 | 0.01765525 | pectin lyase-like superfamily protein; POLYGALACTURONASE LIKE 1 (PGL1) |
| AT1G15510 | -2.60699422 | 2.02E-23 | encodes a pentatricopeptide repeat protein required for chloroplast transcript accD; EARLY CHLOROPLAST BIOGENESIS2 (ECB2) |
| AT1G49475 | -2.58080725 | 0.00625077 | AP2/B3-like transcriptional factor family protein |
| **Time point 12 h** | |  |  |
| ATG | log_2_ FC | p-value | short description |
| AT1G35255 | -15.3928213 | 9.53E-05 | transmembrane protein |
| AT5G20630 | -10.3857629 | 3.98E-09 | encodes a germin-like protein; GERMIN 3 (GER3) |
| AT4G04840 | -4.89398854 | 5.74E-11 | methionine sulfoxide reductase B6 (ATMSRB6) |
| AT5G37260 | -4.59655097 | 2.43E-106 | encodes a MYB family transcription factor Circadian 1 (CIR1); REVEILLE 2 (RVE2) |
| AT5G49330 | -4.38086126 | 8.19E-21 | member of the R2R3 factor gene family; ARABIDOPSIS MYB DOMAIN PROTEIN 111 (ATMYB111) |
| AT1G31690 | -4.36512038 | 3.58E-05 | copper amine oxidase family protein |
| AT1G65060 | -4.33339213 | 2.66E-15 | encodes an isoform of 4-coumarate:CoA ligase (4CL); 4-COUMARATE:COA LIGASE 3 (4CL3) |
| AT1G07167 | -4.33326709 | 0.0023657 | none; long_noncoding_rna |
| AT1G78990 | -4.33226331 | 0.01240497 | HXXXD-type acyl-transferase family protein |
| AT5G13930 | -4.33177305 | 2.09E-07 | encodes chalcone synthase (CHS); TRANSPARENT TESTA 4 (TT4);D127CHALCONE SYNTHASE (CHS) |
| AT3G61920 | -4.31138453 | 0.0024698 | UvrABC system protein C |
| AT5G08640 | -4.20294607 | 1.01E-07 | encodes a flavonol synthase that catalyzes formation of flavonols from dihydroflavonols; FLAVONOL SYNTHASE 1 (FLS1) |
| AT1G49200 | -4.00136931 | 1.64E-14 | RING/U-box superfamily protein |
| AT2G14247 | -3.99342104 | 3.82E-05 | expressed protein |
| AT1G47370 | -3.95262413 | 1.54E-16 | RESPONSE TO THE BACTERIAL TYPE III EFFECTOR PROTEIN HOPBA1 (RBA1) |
| AT5G05270 | -3.95231578 | 1.33E-11 | chalcone-flavanone isomerase family protein; CHALCONE ISOMERASE LIKE (CHIL) |
| AT2G05995 | -3.92767094 | 5.80E-30 | other_RNA |
| AT5G62730 | -3.89355961 | 7.37E-20 | major facilitator superfamily protein |
| AT1G24020 | -3.8902724 | 8.79E-14 | MLP-like protein 423 (MLP423) |
| AT3G07650 | -3.84857331 | 4.52E-146 | CONSTANS-LIKE 9 (COL9); B-BOX DOMAIN PROTEIN 7 (BBX7) |
| AT1G01590 | -3.71503871 | 0.00014502 | FERRIC REDUCTION OXIDASE 1 (FRO1) |

**Supplemental Table S6. Top 20 upregulated genes at different time points after photoperiod stress treatment in *ahk2 ahk3* in comparison to untreated *ahk2 ahk3*.** FC, fold change; p-value according to Bonferroni.

| **Time point 0 h** | |  |  |
| --- | --- | --- | --- |
| ATG | log_2_ FC | p-value | short description |
| AT3G07820 | 16.5051176 | 0.00683985 | Pectin lyase-like superfamily protein |
| AT1G65480 | 7.00006314 | 0.00032475 | FLOWERING LOCUS T (FT) |
| AT1G01060 | 5.36999161 | 1.91E-32 | LATE ELONGATED HYPOCOTYL (LHY) |
| AT5G15830 | 4.57598793 | 1.85E-14 | basic leucine-zipper 3 (bZIP3) |
| AT1G35140 | 3.91553904 | 3.25E-05 | EXL1 is involved in the C-starvation response (EXL1) |
| AT2G46830 | 3.91462872 | 5.47E-35 | CIRCADIAN CLOCK ASSOCIATED 1 (CCA1) |
| AT5G17300 | 3.90331348 | 9.62E-19 | REVEILLE 1 (RVE1) |
| AT5G59990 | 3.83826973 | 1.00E-05 | CCT motif family protein |
| AT1G34580 | 3.78346719 | 0.01021873 | major facilitator superfamily protein |
| AT1G62440 | 3.74674078 | 1.15E-09 | LEUCINE-RICH REPEAT/EXTENSIN 2 (LRX2) |
| AT2G20670 | 3.59668056 | 1.83E-05 | sugar phosphate exchanger, putative (DUF506) |
| AT5G13930 | 3.58523144 | 0.00039508 | encodes chalcone synthase (CHS), a key enzyme involved in the biosynthesis of flavonoids; TRANSPARENT TESTA 4 (TT4);D127CHALCONE SYNTHASE (CHS) |
| AT1G70260 | 3.56293576 | 0.00763585 | encodes an endoplasmic reticulum (ER)-localized nodulin MtN21-like transporter family protein; USUALLY MULTIPLE ACIDS MOVE IN AND OUT TRANSPORTERS 36 (UMAMIT36) |
| AT3G22840 | 3.55200775 | 3.18E-09 | encodes an early light-inducible protein; EARLY LIGHT-INDUCABLE PROTEIN (ELIP1) |
| AT1G69490 | 3.50400622 | 0.00337261 | encodes a member of the NAC transcription factor gene family; ARABIDOPSIS NAC DOMAIN CONTAINING PROTEIN 29 (ANAC029) |
| AT3G49570 | 3.2934664 | 1.12E-12 | response to low sulfur 3 (LSU3) |
| AT2G15880 | 3.25739505 | 0.00011471 | pollen expressed protein required for pollen tube growth; LEUCINE-RICH REPEAT/EXTENSIN 10 (LRX10) |
| AT2G21220 | 3.24505093 | 0.0008296 | SAUR-like auxin-responsive protein family; SMALL AUXIN UPREGULATED RNA 12 (SAUR12) |
| AT3G55980 | 3.23776601 | 5.80E-11 | salt-inducible zinc finger 1 (SZF1) |
| AT2G41250 | 3.18885436 | 4.89E-28 | Haloacid dehalogenase-like hydrolase (HAD) superfamily protein |
| **Time point 4 h** | |  |  |
| ATG | log_2_ FC | p-value | short description |
| AT4G31950 | 10.9483344 | 1.59E-06 | member of CYP82C (CYP82C3) |
| AT1G56250 | 10.6372419 | 6.18E-20 | encodes an F-box protein that can functionally replace VirF, regulating levels of the VirE2 and VIP1 proteins via a VBF-containing SCF complex;PHLOEM PROTEIN 2-B14 (PP2-B14) |
| AT5G51060 | 10.3885031 | 2.59E-16 | RHD2 is required for normal root hair elongation; ROOT HAIR DEFECTIVE 2 (RHD2); RESPIRATORY BURST OXIDASE HOMOLOG C (RBOHC) |
| AT3G02840 | 9.88000436 | 2.43E-29 | ARM repeat superfamily protein |
| AT1G14540 | 9.6617861 | 3.78E-18 | peroxidase superfamily protein; PEROXIDASE 4 (PER4) |
| AT2G37430 | 9.59487894 | 6.60E-18 | ZINC FINGER OF ARABIDOPSIS THALIANA 11 (ZAT11) |
| AT1G47130 | 9.49753441 | 3.16E-06 | hypothetical protein |
| AT3G43250 | 9.2898999 | 1.00E-06 | coiled-coil protein (DUF572) |
| AT4G30280 | 9.22652095 | 1.45E-20 | XYLOGLUCAN ENDOTRANSGLUCOSYLASE/HYDROLASE 18 (XTH18) |
| AT2G23270 | 9.09713776 | 2.74E-06 | transmembrane protein |
| AT1G30370 | 8.93789566 | 4.26E-16 | encodes a mitochondria-localized class III phospholipase A1 that plays a role in seed viability; DAD1-LIKE ACYLHYDROLASE (DLAH) |
| AT4G25810 | 8.92167521 | 3.32E-30 | xyloglucan endotransglycosylase-related protein (XTR6) |
| AT1G56240 | 8.86274751 | 4.94E-15 | phloem protein 2-B13; PHLOEM PROTEIN 2-B13 (PP2-B13) |
| AT5G66890 | 8.84563302 | 1.97E-07 | RPW8 -CNL gene; N REQUIREMENT GENE 1.3 (NRG1.3) |
| AT5G42380 | 8.79044975 | 8.58E-23 | calmodulin like 37 (CML37) |
| AT1G80820 | 8.75302146 | 9.41E-10 | encodes an cinnamoyl CoA reductase isoform; CINNAMOYL COA REDUCTASE (CCR2) |
| AT1G06135 | 8.67503484 | 1.71E-05 | transmembrane protein |
| AT2G02320 | 8.66369662 | 0.00012369 | phloem protein 2-B7; PHLOEM PROTEIN 2-B7 (PP2-B7) |
| AT1G68765 | 8.57311764 | 3.40E-06 | INFLORESCENCE DEFICIENT IN ABSCISSION (IDA) |
| AT1G53620 | 8.55119655 | 1.41E-05 | transmembrane protein |

**Supplemental Table S6 (*continued*). Top 20 upregulated genes at different time points after photoperiod stress treatment in *ahk2 ahk3* in comparison to untreated *ahk2 ahk3*.**

| **Time point 6 h** | |  |  |
| --- | --- | --- | --- |
| ATG | log_2_ FC | p-value | short description |
| AT5G48140 | 28.1608348 | 5.72E-07 | pectin lyase-like superfamily protein |
| AT3G07820 | 17.0596238 | 0.00305495 | pectin lyase-like superfamily protein |
| AT3G01270 | 15.5995993 | 0.00063214 | pectate lyase family protein |
| AT1G05680 | 13.204715 | 4.07E-19 | encodes a UDP-glucosyltransferase; URIDINE DIPHOSPHATE GLYCOSYLTRANSFERASE 74E2 (UGT74E2) |
| AT2G47520 | 12.9730554 | 3.40E-12 | ETHYLENE RESPONSE FACTOR 71 (ERF71); HYPOXIA RESPONSIVE ERF (ETHYLENE RESPONSE FACTOR) 2 (HRE2) |
| AT1G17180 | 12.9630552 | 5.30E-08 | GLUTATHIONE S-TRANSFERASE TAU 25 (GSTU25) |
| AT5G67080 | 12.2841113 | 1.70E-10 | member of MEKK subfamily; MITOGEN-ACTIVATED PROTEIN KINASE KINASE KINASE 19 (MAPKKK19) |
| AT5G66890 | 12.0753742 | 5.25E-13 | RPW8 -CNL gene; N REQUIREMENT GENE 1.3 (NRG1.3) |
| AT5G55150 | 11.7759657 | 5.19E-08 | F-box SKIP23-like protein (DUF295); FBOX/DUF295-RELATED 2 (ATFDR2) |
| AT5G51060 | 11.6476046 | 2.82E-24 | ROOT HAIR DEFECTIVE 2 (RHD2); RESPIRATORY BURST OXIDASE HOMOLOG C (RBOHC) |
| AT2G47550 | 11.6387091 | 7.67E-10 | plant invertase/pectin methylesterase inhibitor superfamily |
| AT4G08770 | 11.4943133 | 6.65E-07 | PEROXIDASE 37 (Prx37) |
| AT5G01380 | 11.4428081 | 4.38E-21 | homeodomain-like superfamily protein |
| AT5G37490 | 11.4389467 | 2.33E-14 | ARM repeat superfamily protein |
| AT5G36970 | 11.3885139 | 5.21E-07 | NDR1/HIN1-LIKE 25 (NHL25) |
| AT4G31950 | 11.3285781 | 3.84E-07 | member of CYP82C; CYTOCHROME P450, FAMILY 82, SUBFAMILY C, POLYPEPTIDE 3 (CYP82C3) |
| AT3G25250 | 11.2582027 | 2.62E-28 | arabidopsis protein kinase; AGC2 KINASE 1 (AGC2-1); OXIDATIVE SIGNAL-INDUCIBLE1 (OXI1) |
| AT3G43250 | 11.2386968 | 1.77E-12 | coiled-coil protein (DUF572) |
| AT2G15490 | 11.1443601 | 8.24E-07 | UDP-glycosyltransferase 73B4; UDP-GLYCOSYLTRANSFERASE 73B4 (UGT73B4) |
| AT4G30280 | 10.9725774 | 1.03E-29 | XYLOGLUCAN ENDOTRANSGLUCOSYLASE/HYDROLASE 18 (XTH18) |
| **Time point 12 h** | |  |  |
| ATG | log_2_ FC | p-value | short description |
| AT5G48140 | 28.904847 | 1.18E-07 | pectin lyase-like superfamily proteinp |
| AT2G42560 | 19.1500032 | 7.84E-09 | LATE EMBRYOGENESIS ABUNDANT 25 (LEA25) |
| AT3G01420 | 13.9174363 | 0.00032175 | PLANT ALPHA DIOXYGENASE 1 (PADOX-1) |
| AT1G15380 | 10.9407994 | 0.00649811 | GLYOXYLASE I 4 (GLYI4) |
| AT1G01460 | 10.2186989 | 1.46E-09 | type I phosphatidylinositol-4-phosphate 5-kinase, subfamily A (PIPK11) |
| AT2G32460 | 9.83519095 | 1.56E-08 | member of the R2R3 factor gene family; MYB DOMAIN PROTEIN 101 (MYB101); ABNORMAL SHOOT 7 (ABS7); ABC TRANSPORTER OF THE MITOCHONDRION 1 (ATM1) |
| AT3G61930 | 9.8294985 | 0.00057798 | hypothetical protein |
| AT1G69880 | 9.71986331 | 0.00012811 | thioredoxin H-type 8; THIOREDOXIN H-TYPE 8 (TH8) |
| AT1G70440 | 9.34110097 | 1.19E-28 | SIMILAR TO RCD ONE 3 (SRO3) |
| AT3G60140 | 9.17120419 | 0.00523602 | DARK INDUCIBLE 2 (DIN2) SENESCENCE-RELATED GENE 2 (SRG2);BETA GLUCOSIDASE 30 (BGLU30) |
| AT3G02375 | 9.11332244 | 9.10E-10 | novel transcribed region; detected in light-grown seedling |
| AT5G35380 | 9.01081052 | 0.00017488 | kinase with adenine nucleotide alpha hydrolases-like domain-containing protein |
| AT3G03670 | 9.00659716 | 0.00643281 | peroxidase superfamily protein |
| AT1G18860 | 8.97365884 | 0.00514896 | WRKY DNA-BINDING PROTEIN 61 (WRKY61) |
| AT5G14602 | 8.96842278 | 8.16E-06 | methyltransferase-like protein |
| AT5G06839 | 8.9176204 | 6.92E-23 | bZIP transcription factor family protein; TGACG (TGA) MOTIF-BINDING PROTEIN 10 (TGA10) (bZIP65) |
| AT4G19000 | 8.90074475 | 2.95E-05 | the C-terminal portion of this protein has homology to the C-termini of the IWS1 (Interacts With Spt6) proteins found in yeast and humans (IWS2) |
| AT4G28940 | 8.90028986 | 1.73E-10 | phosphorylase superfamily protein |
| AT5G50760 | 8.69818074 | 1.59E-18 | SMALL AUXIN UPREGULATED RNA 55 (SAUR55) |
| AT1G67220 | 8.65187451 | 7.23E-12 | HISTONE ACETYLTRANSFERASE OF THE CBP FAMILY 2 (HAC2) |

**Supplemental Table S7. Top 20 downregulated genes at different time points after photoperiod stress treatment in *ahk2 ahk3* in comparison to untreated *ahk2 ahk3*.** FC, fold change; p-value according to Bonferroni.

| **Time point 0 h** | |  |  |
| --- | --- | --- | --- |
| ATG | log_2_ FC | p-value | short description |
| AT1G70640 | -3.23292842 | 8.11E-10 | octicosapeptide/Phox/Bem1p (PB1) domain-containing protein |
| AT4G36570 | -3.19292817 | 1.86E-08 | RAD-like 3 (RL3) |
| AT5G15960 | -2.90423291 | 0.04167381 | cold and ABA inducible protein kin1, possibly functions as an anti-freeze protein (KIN1) |
| AT1G64220 | -2.79443786 | 0.03188318 | translocase of outer membrane 7 kDa subunit 2 (TOM7-2) |
| AT5G60730 | -2.73355191 | 2.85E-07 | GUIDED ENTRY OF TAIL-ANCHORED PROTEIN 3C (ATGET3C) |
| AT1G51380 | -2.67828376 | 1.39E-10 | DEA(D/H)-box RNA helicase family protein |
| AT3G22235 | -2.64327846 | 0.00122588 | cysteine-rich TM module stress tolerance protein (ATHCYSTM8) |
| AT3G01960 | -2.61784665 | 0.00061932 | hypothetical protein |
| AT5G11590 | -2.56053957 | 2.91E-09 | encodes a member of the DREB subfamily A-4 of ERF/AP2 transcription factor family; TINY2 (TINY2) |
| AT2G42540 | -2.45618529 | 0.02662268 | COLD-REGULATED 15A (COR15A) |
| AT4G26150 | -2.42376521 | 6.06E-07 | CYTOKININ-RESPONSIVE GATA FACTOR 1 (CGA1) |
| AT5G59670 | -2.3932599 | 4.32E-09 | Leucine-rich repeat protein kinase family protein |
| AT3G51400 | -2.28969113 | 2.76E-09 | hypothetical protein (DUF241) |
| AT1G68550 | -2.27261828 | 2.04E-26 | CYTOKININ RESPONSE FACTOR 10 (CRF10) |
| AT2G05995 | -2.2149446 | 2.20E-09 | other_RNA |
| AT5G59662 | -2.20836626 | 2.41E-05 | natural antisense transcript overlaps with AT5G59660 |
| AT5G14580 | -2.14870838 | 9.51E-17 | polyribonucleotide nucleotidyltransferase |
| AT2G40080 | -2.13249565 | 1.08E-26 | EARLY FLOWERING 4 (ELF4) |
| AT5G06550 | -2.129209 | 5.85E-12 | JUMONJI DOMAIN-CONTAINING PROTEIN 22 (JMJ22) |
| AT2G42530 | -2.1158961 | 2.48E-09 | COLD REGULATED 15B (COR15B) |
| AT1G16830 | -2.11328006 | 5.02E-05 | pentatricopeptide repeat (PPR) superfamily protein |
| AT2G47880 | -2.10127049 | 0.00045046 | encodes a member of the CC-type glutaredoxin (ROXY) family; CEP DOWNSTREAM 2 (CEPD2) (ROXY9) |
| AT4G16750 | -2.09787396 | 8.04E-11 | ETHYLENE-RESPONSIVE TRANSCRIPTION FACTOR 39 (ERF39) |
| AT3G05660 | -2.09681737 | 3.44E-06 | receptor like protein 33 (RLP33) |
| AT3G03060 | -2.09322123 | 4.03E-08 | P-loop containing nucleoside triphosphate hydrolases superfamily protein |
| AT3G59670 | -2.0915183 | 1.47E-11 | elongation factor |
| **Time point 4 h** | |  |  |
| ATG | log_2_ FC | p-value | short description |
| AT2G47040 | -35.263979 | 5.36E-30 | VANGUARD1 (VGD1) |
| AT3G07820 | -15.9896234 | 0.01602478 | pectin lyase-like superfamily protein |
| AT1G60590 | -4.63999795 | 3.35E-08 | pectin lyase-like superfamily |
| AT1G52342 | -4.40148966 | 1.96E-23 | hypothetical protein |
| AT1G31173 | -4.25564195 | 2.56E-06 | MICRORNA167D (MIR167D) |
| AT2G41240 | -4.22541142 | 0.00182195 | BASIC HELIX-LOOP-HELIX PROTEIN 100 (BHLH100) |
| AT5G03350 | -4.18500334 | 0.03749811 | SA-INDUCED LEGUME LECTIN-LIKE PROTEIN 1 (SAI-LLP1) |
| AT5G56100 | -3.91909306 | 6.30E-22 | glycine-rich protein / oleosin |
| AT5G04150 | -3.89029087 | 0.00012022 | BASIC HELIX-LOOP-HELIX PROTEIN 101 (BHLH101) |
| AT4G25780 | -3.75612341 | 2.40E-05 | CAP (Cysteine-rich secretory proteins, Antigen 5, and Pathogenesis-related 1 protein) superfamily protein (ATCAPE2) |
| AT4G15990 | -3.42762891 | 1.15E-07 | hypothetical protein |
| AT3G17640 | -3.23053832 | 4.76E-12 | leucine-rich repeat (LRR) family protein |
| AT3G58070 | -3.22231623 | 3.30E-12 | GLABROUS INFLORESCENCE STEMS (GIS) |
| AT2G14247 | -3.12777321 | 0.03747838 | expressed protein |
| AT2G05160 | -3.1195057 | 3.75E-20 | CCCH-type zinc fingerfamily protein with RNA-binding domain-containing protein |
| AT1G78930 | -2.98719812 | 1.95E-12 | mitochondrial transcription termination factor family protein |
| AT1G32583 | -2.9346286 | 0.00010898 | a microRNA MIR400 is derived from the first intron of At1g32583 in the 5'UTR |
| AT4G26530 | -2.91941301 | 1.23E-08 | FRUCTOSE-BISPHOSPHATE ALDOLASE 5 (FBA5) |
| AT5G18404 | -2.91824467 | 0.00035771 | this gene encodes a small protein and has either evidence of transcription or purifying selection |
| AT2G46660 | -2.8996421 | 3.50E-15 | CYTOCHROME P450, FAMILY 78, SUBFAMILY A, POLYPEPTIDE 6 (CYP78A6) |

**Supplemental Table S7 (*continued*). Top 20 downregulated genes at different time points after photoperiod stress treatment in *ahk2 ahk3* in comparison to untreated *ahk2 ahk3*.**

| **Time point 6 h** | |  |  |
| --- | --- | --- | --- |
| ATG | log_2_ FC | p-value | short description |
| AT1G31173 | -6.94557448 | 0.00024036 | encodes a microRNA that targets ARF family members ARF6 and ARF8; MICRORNA167D (MIR167D) |
| AT3G28270 | -6.86399615 | 6.90E-19 | AT14A-LIKE1 (AFL1) |
| AT4G36570 | -6.15559525 | 6.97E-12 | RAD-like 3 (RL3) |
| AT5G59662 | -6.12602623 | 0.00987881 | natural antisense transcript overlaps with AT5G59660 |
| AT1G60590 | -5.56607502 | 6.71E-10 | pectin lyase-like superfamily protein |
| AT5G03350 | -5.54155466 | 0.00359236 | SA-INDUCED LEGUME LECTIN-LIKE PROTEIN 1 (SAI-LLP1) |
| AT5G20630 | -5.34116538 | 3.29E-07 | encodes a germin-like protein; GERMIN 3 (GER3) |
| AT3G61510 | -5.26297066 | 4.83E-05 | ARABIDOPSIS THALIANA 1-AMINOCYCLOPROPANE-1-CARBOXYLATE SYNTHASE 1 (ACS1) |
| AT4G26530 | -5.25917244 | 5.62E-33 | FRUCTOSE-BISPHOSPHATE ALDOLASE 5 (FBA5) |
| AT2G05160 | -5.13670886 | 1.59E-32 | CCCH-type zinc fingerfamily protein with RNA-binding domain-containing protein |
| AT4G12320 | -4.96605016 | 5.02E-33 | CYTOCHROME P450, FAMILY 706, SUBFAMILY A, POLYPEPTIDE 6 (CYP706A6) |
| AT2G14247 | -4.92048452 | 8.42E-09 | expressed protein |
| AT2G41240 | -4.77327828 | 3.93E-05 | BASIC HELIX-LOOP-HELIX PROTEIN 100 (BHLH100) |
| AT4G12830 | -4.76034869 | 7.48E-43 | alpha/beta-Hydrolases superfamily protein |
| AT1G29510 | -4.73159224 | 0.01661831 | SMALL AUXIN UPREGULATED RNA 67 (SAUR67) |
| AT4G12970 | -4.6919243 | 5.58E-17 | STOMAGEN (STOMAGEN); (ATEPFL9);EPIDERMAL PATTERNING FACTOR LIKE-9 (EPFL9) |
| AT3G58070 | -4.6472497 | 2.91E-17 | GLABROUS INFLORESCENCE STEMS (GIS) |
| AT3G62550 | -4.62519048 | 6.40E-46 | adenine nucleotide alpha hydrolases-like superfamily protein |
| AT1G30250 | -4.59403941 | 1.05E-29 | hypothetical protein |
| AT3G11110 | -4.45853683 | 2.32E-34 | RING/U-box superfamily protein |
| **Time point 12 h** | |  |  |
| ATG | log_2_ FC | p-value | short description |
| AT1G35255 | -16.325313 | 7.50E-07 | transmembrane protein |
| AT5G18020 | -9.57531321 | 3.38E-10 | SAUR-like auxin-responsive protein family; SMALL AUXIN UP RNA 20 (SAUR20) |
| AT1G13609 | -9.05544546 | 1.51E-12 | encodes a defensin-like (DEFL) family protein |
| AT5G20630 | -8.68667406 | 2.75E-14 | GERMIN 3 (GER3) |
| AT3G10150 | -8.51885364 | 9.63E-08 | PURPLE ACID PHOSPHATASE 16 (PAP16) |
| AT5G04530 | -8.48222985 | 1.90E-06 | 3-KETOACYL-COA SYNTHASE 19 (KCS19) |
| AT5G18030 | -8.45323061 | 3.37E-07 | SMALL AUXIN UP RNA 21 (SAUR21) |
| AT1G29450 | -8.24544546 | 1.64E-09 | SMALL AUXIN UPREGULATED RNA 64 (SAUR64) |
| AT3G17640 | -8.20371726 | 6.75E-07 | Leucine-rich repeat (LRR) family protein |
| AT2G18120 | -8.1722531 | 5.65E-07 | SHI-RELATED SEQUENCE 4 (SRS4) |
| AT3G03840 | -8.16120768 | 2.62E-06 | SMALL AUXIN UP RNA 27 (SAUR27) |
| AT5G50800 | -8.16106505 | 2.72E-06 | encodes a member of the SWEET sucrose efflux transporter family proteins (SWEET13); RUPTURED POLLEN GRAIN 2 (RPG2) |
| AT3G56970 | -7.80088264 | 9.03E-06 | BASIC HELIX-LOOP-HELIX 38 (BHLH38) |
| AT1G43605 | -7.74420769 | 1.20E-05 | hypothetical protein |
| AT3G28070 | -7.70926547 | 1.04E-42 | nodulin MtN21-like transporter family protein; USUALLY MULTIPLE ACIDS MOVE IN AND OUT TRANSPORTERS 46 (UMAMIT46) |
| AT5G46690 | -7.69967552 | 4.57E-13 | beta HLH protein 71; BETA HLH PROTEIN 71 (bHLH071) |
| AT2G41240 | -7.618768 | 8.68E-14 | BASIC HELIX-LOOP-HELIX PROTEIN 100 (BHLH100) |
| AT3G63450 | -7.49485149 | 9.25E-06 | RNA-binding (RRM/RBD/RNP motifs) |
| AT5G47610 | -7.49091879 | 7.65E-34 | RING/U-box superfamily protein |
| AT1G24020 | -7.41722087 | 4.18E-05 | MLP-LIKE PROTEIN 423 (MLP423) |

**Supplemental Table 8. Top 20 upregulated genes at different time points after photoperiod stress treatment in *cca1 lhy* in comparison to untreated *cca1 lhy*.** FC, fold change; p-value according to Bonferroni.

| **Time point 0 h** | |  |  |
| --- | --- | --- | --- |
| ATG | log_2_ FC | p-value | short description |
| AT2G47040 | 39.4707407 | 3.24E-40 | VANGUARD1 (VGD1) |
| AT5G48140 | 31.6093898 | 1.58E-10 | pectin lyase-like superfamily protein |
| AT1G35255 | 17.1685735 | 1.16E-07 | transmembrane protein |
| AT5G45820 | 6.4229615 | 0.00061093 | CBL-INTERACTING PROTEIN KINASE 20 (CIPK20); SNF1-RELATED PROTEIN KINASE 3.6 (SnRK3.6); PROTEIN KINASE 18 (PKS18) |
| AT2G46830 | 5.9191757 | 0.0005438 | CIRCADIAN CLOCK ASSOCIATED 1 (CCA1) |
| AT1G70260 | 5.51846343 | 4.03E-14 | USUALLY MULTIPLE ACIDS MOVE IN AND OUT TRANSPORTERS 36 (UMAMIT36) |
| AT5G03545 | 4.3267951 | 0.00505321 | INDUCED BY PI STARVATION 2 (ATIPS2) |
| AT3G08040 | 4.17111691 | 0.00200678 | FERRIC REDUCTASE DEFECTIVE 3 (FRD3) |
| AT3G02040 | 3.53135942 | 6.48E-06 | SENESCENCE-RELATED GENE 3 (SRG3); GLYCEROPHOSPHODIESTER PHOSPHODIESTERASE 1 (GDPD1) |
| AT1G29920 | 3.48162798 | 6.41E-11 | CHLOROPHYLL A/B-BINDING PROTEIN 2 (CAB2) |
| AT4G38340 | 3.35943507 | 2.78E-08 | NIN-LIKE PROTEIN 3 (NLP3) |
| AT3G24460 | 3.30468534 | 8.08E-34 | serinc-domain containing serine and sphingolipid biosynthesis protein |
| AT5G15830 | 3.25809148 | 2.01E-09 | BASIC LEUCINE-ZIPPER 3 (bZIP3) |
| AT3G09600 | 3.19877343 | 1.63E-41 | REVEILLE 8 (RVE8) |
| AT4G20820 | 3.12200754 | 2.27E-15 | FAD-binding Berberine family protein (ATBBE18) |
| AT1G01520 | 3.0732288 | 8.04E-08 | ALTERED SEED GERMINATION 4 (ASG4); REVEILLE 3 (REV3) |
| AT5G59340 | 3.04660506 | 3.45E-12 | WUSCHEL RELATED HOMEOBOX 2 (WOX2) |
| AT5G17300 | 3.03635302 | 5.28E-19 | REVEILLE 1 (RVE1) |
| AT3G54500 | 3.02353741 | 1.47E-31 | NIGHT LIGHT-INDUCIBLE AND CLOCK-REGULATED 2 (LNK2) |
| AT1G21910 | 3.00260839 | 2.75E-07 | DEHYDRATION RESPONSE ELEMENT-BINDING PROTEIN 26 (DREB26) |
| **Time point 4 h** | |  |  |
| ATG | log_2_ FC | p-value | short description |
| AT1G35255 | 17.4611062 | 5.01E-08 | transmembrane protein |
| AT3G01270 | 16.8465159 | 7.10E-06 | pectate lyase family protein |
| AT1G08860 | 10.4975255 | 1.66E-07 | BONZAI 3 (BON3) |
| AT5G28610 | 8.79833046 | 1.73E-05 | LOW protein: ATP-dependent RNA helicase DRS1-like protein |
| AT5G38700 | 8.51444612 | 0.00048886 | cotton fiber protein |
| AT1G65390 | 7.81417327 | 1.83E-17 | PHLOEM PROTEIN 2 A5 (PP2-A5) |
| AT4G30430 | 7.78636844 | 8.71E-08 | TETRASPANIN9 (TET9) |
| AT4G30280 | 7.75733593 | 8.13E-14 | XYLOGLUCAN ENDOTRANSGLUCOSYLASE/HYDROLASE 18 (XTH18) |
| AT1G14540 | 7.74268919 | 2.62E-11 | peroxidase superfamily protein; PEROXIDASE 4 (PER4) |
| AT1G56250 | 7.4334234 | 1.11E-11 | PHLOEM PROTEIN 2-B14 (PP2-B14) |
| AT2G23270 | 7.42475907 | 0.00379852 | transmembrane protein |
| AT5G56960 | 7.36293833 | 4.83E-16 | basic helix-loop-helix (bHLH) DNA-binding family protein |
| AT1G73550 | 7.27215647 | 0.00408163 | encodes a Protease inhibitor/seed storage/LTP family protein |
| AT4G24110 | 7.1666458 | 3.11E-15 | NADP-specific glutamate dehydrogenase |
| AT1G70260 | 7.16427762 | 6.59E-12 | USUALLY MULTIPLE ACIDS MOVE IN AND OUT TRANSPORTERS 36 (UMAMIT36) |
| AT1G56240 | 7.09882452 | 3.73E-08 | phloem protein 2-B13; PHLOEM PROTEIN 2-B13 (PP2-B13) |
| AT1G56060 | 7.08880861 | 1.79E-13 | CYSTEINE-RICH TRANSMEMBRANE MODULE 3 (ATHCYSTM3) |
| AT1G65310 | 7.08503138 | 0.01590895 | XYLOGLUCAN ENDOTRANSGLUCOSYLASE/HYDROLASE 17 (XTH17) |
| AT2G37430 | 7.00866234 | 1.64E-13 | ZINC FINGER OF ARABIDOPSIS THALIANA 11 (ZAT11) |
| AT1G16420 | 6.82398088 | 1.22E-10 | METACASPASE 8 (MC8) |

**Supplemental Table S8 (*continued*). Top 20 upregulated genes at different time points after photoperiod stress treatment in *cca1 lhy* in comparison to untreated *cca1 lhy*.**

| **Time point 6 h** | |  |  |
| --- | --- | --- | --- |
| ATG | log_2_ FC | p-value | short description |
| AT1G66550 | 18.9443672 | 4.12E-11 | WRKY DNA-BINDING PROTEIN 67 (WRKY67) |
| AT1G17180 | 11.3969235 | 1.44E-05 | GLUTATHIONE S-TRANSFERASE TAU 25 (ATGSTU25) |
| AT5G66890 | 11.2210271 | 6.99E-11 | N REQUIREMENT GENE 1.3 (NRG1.3) N REQUIREMENT GENE 1.3 (NRG1.3) |
| AT5G24640 | 10.4418814 | 1.85E-07 | hypothetical protein |
| AT4G30280 | 10.2894844 | 8.50E-26 | XYLOGLUCAN ENDOTRANSGLUCOSYLASE/HYDROLASE 18 (ATXTH18) |
| AT1G14540 | 10.0480638 | 3.00E-18 | PEROXIDASE 4 (PER4) |
| AT1G56060 | 10.0440685 | 1.13E-28 | CYSTEINE-RICH TRANSMEMBRANE MODULE 3 (ATHCYSTM3) |
| AT3G43250 | 9.8141477 | 4.43E-05 | coiled-coil protein (DUF572) |
| AT5G42380 | 9.72602175 | 2.56E-28 | CALMODULIN LIKE 37 (CML37) |
| AT5G38700 | 9.62277173 | 5.61E-06 | cotton fiber protein |
| AT3G25250 | 9.61893558 | 1.58E-21 | AGC2 KINASE 1 (AGC2-1);OXIDATIVE SIGNAL-INDUCIBLE1 (OXI1) |
| AT1G56250 | 9.60721291 | 9.12E-16 | PHLOEM PROTEIN 2-B14 (PP2-B14) |
| AT5G55150 | 9.55392985 | 2.44E-05 | FBOX/DUF295-RELATED 2 (ATFDR2) |
| AT3G46080 | 9.53167792 | 5.35E-19 | C2H2-type zinc finger family protein |
| AT5G01380 | 9.39628545 | 2.70E-16 | homeodomain-like superfamily protein |
| AT1G32350 | 9.23961479 | 2.05E-08 | alternative oxidase 1D; ALTERNATIVE OXIDASE 1D (AOX1D) |
| AT2G02010 | 9.19771955 | 2.77E-25 | GLUTAMATE DECARBOXYLASE 4 (GAD4) GLUTAMATE DECARBOXYLASE 4 (GAD4) |
| AT1G13520 | 9.17123183 | 1.29E-07 | hypothetical protein (DUF1262) |
| AT3G62760 | 9.15851226 | 1.28E-05 | encodes glutathione transferase belonging to the phi class of GSTs (ATGSTF13) |
| AT1G48640 | 9.04460914 | 3.08E-07 | transmembrane amino acid transporter family protein |
| **Time point 12 h** | |  |  |
| ATG | log_2_ FC | p-value | short description |
| AT1G15540 | 8.22322524 | 1.39E-05 | 2-oxoglutarate-dependent dioxygenase-like protein |
| AT4G34470 | 8.20382836 | 7.29E-07 | SKP1-LIKE 12 (SK12) |
| AT2G14620 | 7.81989246 | 1.67E-21 | XYLOGLUCAN ENDOTRANSGLUCOSYLASE/HYDROLASE 10 (XTH10) |
| AT1G74080 | 7.59192993 | 1.88E-10 | MYB DOMAIN PROTEIN 122 (MYB122) |
| AT4G34210 | 7.50445353 | 1.60E-05 | one of 20 SKP1 homologs in Arabidopsis. Protein is most similar to ASK12 and RNAi lines show defects in stamen development; SKP1-LIKE 11 (SK11) |
| AT3G11340 | 7.35370106 | 0.00104297 | UDP-DEPENDENT GLYCOSYLTRANSFERASE 76B1 (UGT76B1) |
| AT1G70440 | 7.3350161 | 5.70E-19 | SIMILAR TO RCD ONE 3 (SRO3) |
| AT5G49690 | 7.27426013 | 2.26E-11 | UDP-Glycosyltransferase superfamily protein |
| AT5G24206 | 7.22359598 | 2.60E-33 | other_RNA |
| AT4G16260 | 7.18162663 | 2.90E-11 | encodes a putative beta-1,3-endoglucanase that interacts with the 30C02 cyst nematode effector. |
| AT3G03260 | 7.02242738 | 0.00659355 | HOMEODOMAIN GLABROUS 8 (HDG8) HOMEODOMAIN GLABROUS 8 (HDG8) |
| AT1G49900 | 6.99658735 | 2.60E-18 | C2H2 type zinc finger transcription factor family |
| AT1G01460 | 6.9767701 | 6.73E-06 | Type I phosphatidylinositol-4-phosphate 5-kinase, subfamily A (PIPK11) |
| AT2G46750 | 6.97648446 | 0.00229549 | encodes a homolog of rat L-gulono-1,4-lactone (L-GulL) oxidase that is involved in the biosynthesis of L-ascorbic acid; L -GULONO-1,4-LACTONE ( L -GULL) OXIDASE 2 (GULLO2) |
| AT3G44830 | 6.86624805 | 4.61E-06 | lecithin:cholesterol acyltransferase family protein |
| AT1G65970 | 6.73679855 | 4.85E-06 | THIOREDOXIN-DEPENDENT PEROXIDASE 2 (TPX2) |
| AT1G60750 | 6.6922404 | 0.00884061 | NAD(P)-linked oxidoreductase superfamily protein |
| AT1G62490 | 6.66562259 | 0.00026817 | mitochondrial transcription termination factor family protein |
| AT4G08770 | 6.64958605 | 0.00539371 | PEROXIDASE 37 (Prx37) |
| AT3G03650 | 6.61313718 | 0.00999608 | EMBRYO SAC DEVELOPMENT ARREST 5 (EDA5) |

**Supplemental Table S9. Top 20 downregulated genes at different time points after photoperiod stress treatment in *cca1 lhy* in comparison to untreated *cca1 lhy*.** FC, fold change; p-value according to Bonferroni.

| **Time point 0 h** | |  |  |
| --- | --- | --- | --- |
| ATG | log_2_ FC | p-value | short description |
| AT1G66550 | -23.9823092 | 2.57E-24 | WRKY DNA-BINDING PROTEIN 67 (WRKY67) |
| AT4G17470 | -3.60181212 | 0.0396289 | alpha/beta-Hydrolases superfamily protein |
| AT1G52030 | -3.35085319 | 0.0121987 | MYROSINASE-BINDING PROTEIN 2 (MBP2) |
| AT3G05660 | -3.12064896 | 1.58E-15 | RECEPTOR LIKE PROTEIN 33 (RLP33) |
| AT3G59710 | -2.84956434 | 0.00036565 | NAD(P)-binding Rossmann-fold superfamily protein |
| AT5G27060 | -2.69514013 | 0.01930247 | receptor like protein 53; RECEPTOR LIKE PROTEIN 53 (RLP53) |
| AT1G24090 | -2.60283209 | 1.22E-12 | RNase H family protein (RNH1C) |
| AT5G60730 | -2.56130333 | 5.51E-06 | GUIDED ENTRY OF TAIL-ANCHORED PROTEINS 3B (GET3C) |
| AT5G15840 | -2.43879865 | 3.65E-06 | CONSTANS (CO); B-BOX DOMAIN PROTEIN 1 (BBX1) |
| AT5G23820 | -2.39997618 | 0.00687216 | MD2-RELATED LIPID RECOGNITION 3 (ML3) |
| AT1G18320 | -2.25136269 | 0.04053292 | mitochondrial import inner membrane translocase subunit Tim17/Tim22/Tim23 family protein |
| AT3G20440 | -2.2396177 | 2.07E-14 | EMBRYO DEFECTIVE 2729 (EMB2729) |
| AT3G49320 | -2.23620081 | 1.96E-15 | metal-dependent protein hydrolase |
| AT4G30650 | -2.12260203 | 2.98E-18 | low temperature and salt responsive protein family |
| AT1G51380 | -2.0798619 | 3.10E-05 | DEA(D/H)-box RNA helicase family protein |
| AT5G15970 | -2.05325852 | 4.10E-13 | COLD-RESPONSIVE 6.6 (COR6.6) |
| AT2G37690 | -2.03577619 | 1.38E-18 | phosphoribosylaminoimidazole carboxylase |
| AT5G66960 | -1.9963076 | 9.70E-07 | prolyl oligopeptidase family protein |
| AT5G59662 | -1.99622155 | 5.40E-07 | natural antisense transcript overlaps with AT5G59660 |
| AT5G06550 | -1.93310231 | 8.10E-09 | JUMONJI DOMAIN-CONTAINING PROTEIN 22 (JMJ22) |
| **Time point 4 h** | |  |  |
| ATG | log_2_ FC | p-value | short description |
| AT1G66550 | -19.3019558 | 2.00E-13 | WRKY DNA-BINDING PROTEIN 67 (WRKY67) |
| AT1G53540 | -8.00698845 | 0.02927744 | HSP20-like chaperones superfamily protein (HSP17.6C) |
| AT4G11911 | -7.73031972 | 0.00118439 | STAY-GREEN-like protein |
| AT4G35160 | -7.40932771 | 0.00076152 | N-ACETYLSEROTONIN O-METHYLTRANSFERASE (ASMT) |
| AT5G24780 | -6.1715481 | 1.07E-05 | VEGETATIVE STORAGE PROTEIN 1 (VSP1) |
| AT5G07010 | -6.15753774 | 1.63E-05 | SULFOTRANSFERASE 2A (ST2A) |
| AT2G39030 | -5.5411476 | 8.08E-06 | N-ACETYLTRANSFERASE ACTIVITY 1 (NATA1) |
| AT4G17470 | -5.51862547 | 1.60E-08 | alpha/beta-Hydrolases superfamily protein |
| AT4G11320 | -5.09230615 | 0.00132041 | papain family cysteine protease; CYSTEINE PROTEASE 2 (CP2) |
| AT5G63450 | -4.82552393 | 0.04059716 | CYTOCHROME P450, FAMILY 94, SUBFAMILY B, POLYPEPTIDE 1 (CYP94B1) |
| AT3G28740 | -4.70929043 | 0.00036255 | CYTOCHROME P450, FAMILY 81, SUBFAMILY D, POLYPEPTIDE 11 (CYP81D11) |
| AT2G24850 | -4.7052898 | 0.00825378 | TYROSINE AMINOTRANSFERASE 3 (TAT3) |
| AT5G24770 | -4.52045175 | 0.0008433 | VEGETATIVE STORAGE PROTEIN 2 (VSP2) |
| AT1G73325 | -4.29138301 | 0.00087474 | Kunitz family trypsin and protease inhibitor protein |
| AT1G54010 | -4.267132 | 0.0004799 | GDSL-LIKE LIPASE 23 (GLL23) |
| AT5G03350 | -4.11551998 | 0.01636523 | SA-INDUCED LEGUME LECTIN-LIKE PROTEIN 1 (SAI-LLP1) |
| AT4G15440 | -3.75545028 | 8.74E-15 | HYDROPEROXIDE LYASE 1 (HPL1) |
| AT3G14280 | -3.65350143 | 0.00423749 | LL-diaminopimelate aminotransferase |
| AT4G15210 | -3.64954813 | 0.01079944 | BETA-AMYLASE 5 (BAM5) |
| AT2G43510 | -3.63096034 | 0.00668202 | ATRYPSIN INHIBITOR PROTEIN 1 (TI1) |

**Supplemental Table S9 (*continued*). Top 20 downregulated genes at different time points after photoperiod stress treatment in *cca1 lhy* in comparison to untreated *cca1 lhy*.**

| **Time point 6 h** | |  |  |
| --- | --- | --- | --- |
| ATG | log_2_ FC | p-value | short description |
| AT3G62170 | -30.8376221 | 0.0149844 | VANGUARD-like protein; VANGUARD 1 HOMOLOG 2 (VGDH2) |
| AT1G07050 | -4.94434871 | 3.85E-55 | FITNESS encodes a protein with a single CCT domain and belongs to the CCT motif family genes (CMF). FITNESS acts upstream JUB1 thereby controlling H2O2 levels. FITNESS (FITNESS) |
| AT1G73325 | -4.70440741 | 0.0012485 | Kunitz family trypsin and protease inhibitor protein |
| AT3G15720 | -4.31024262 | 2.33E-12 | pectin lyase-like superfamily protein |
| AT4G17470 | -4.23071658 | 0.00166344 | alpha/beta-Hydrolases superfamily protein |
| AT1G28330 | -4.0350969 | 4.39E-10 | dormancy-associated protein; DORMANCY-ASSOCIATED PROTEIN-LIKE 1 (DYL1) |
| AT1G56650 | -3.84780919 | 1.67E-05 | PRODUCTION OF ANTHOCYANIN PIGMENT 1 (PAP1) |
| AT4G15440 | -3.82445016 | 7.31E-15 | encodes a hydroperoxide lyase; HYDROPEROXIDE LYASE 1 (HPL1) |
| AT1G30250 | -3.76525235 | 4.70E-23 | hypothetical protein |
| AT3G63210 | -3.71074898 | 3.80E-20 | encodes a novel zinc-finger protein with a proline-rich N-terminus, identical to senescence-associated protein SAG102; MEDIATOR OF ABA-REGULATED DORMANCY 1 (MARD1) |
| AT5G05250 | -3.64505293 | 1.28E-20 | hypothetical protein |
| AT3G62550 | -3.59761423 | 4.35E-27 | adenine nucleotide alpha hydrolases-like superfamily protein |
| AT2G36400 | -3.56428094 | 1.36E-10 | GROWTH-REGULATING FACTOR 3 (GRF3) |
| AT2G43535 | -3.51079038 | 2.77E-16 | encodes a defensin-like (DEFL) family protein |
| AT3G26740 | -3.50318234 | 5.11E-50 | CCR-LIKE (CCL) |
| AT5G61890 | -3.45952865 | 0.00047818 | encodes a member of the ERF (ethylene response factor) subfamily B-4 of ERF/AP2 transcription factor family (ERF114) |
| AT1G22370 | -3.44704509 | 2.58E-18 | UDP-GLUCOSYL TRANSFERASE 85A5 (UGT85A5) |
| AT4G39110 | -3.4168149 | 0.00350018 | BUDDHAS PAPER SEAL 1 (BUPS1) |
| AT5G16410 | -3.37881114 | 0.00034606 | HXXXD-type acyl-transferase family protein |
| AT5G07700 | -3.35412901 | 5.63E-07 | encodes a putative transcription factor (MYB76); MYB DOMAIN PROTEIN 76 (MYB76) |
| **Time point 12 h** | |  |  |
| ATG | log_2_ FC | p-value | short description |
| AT1G35255 | -14.8418469 | 0.00034784 | transmembrane protein |
| AT2G27420 | -7.71999094 | 1.45E-05 | cysteine proteinases superfamily protein |
| AT1G26790 | -6.63998274 | 4.56E-41 | Dof-type zinc finger DNA-binding family protein; CYCLING DOF FACTOR 6 (CDF6) |
| AT3G29590 | -6.61946685 | 0.03129318 | encodes a malonyl-CoA:anthocyanidin 5-O-glucoside-6"-O-malonyltransferase that is coordinately expressed with a epistatic 5-O-anthocyanidin glucosyltransferase (At4g14090) (AT5MAT) |
| AT5G37300 | -6.10120525 | 0.00536128 | encodes a bifunctional enzyme, wax ester synthase (WS) and diacylglycerol acyltransferase (DGAT) (WSD1) |
| AT1G65060 | -5.78373527 | 6.23E-23 | 4-COUMARATE:COA LIGASE 3 (4CL3) |
| AT3G18217 | -5.74669772 | 8.00E-07 | MICRORNA157C (MIR157C) |
| AT1G13609 | -5.58717999 | 1.34E-06 | encodes a defensin-like (DEFL) family protein |
| AT5G59320 | -5.55393753 | 0.00010907 | LIPID TRANSFER PROTEIN 3 (LTP3) |
| AT4G16590 | -5.52458493 | 0.00059126 | CELLULOSE SYNTHASE-LIKE A01 (CSLA01) |
| AT3G28270 | -5.491454 | 2.35E-15 | AT14A-LIKE1 (AFL1) |
| AT1G02205 | -5.36476789 | 2.07E-06 | ECERIFERUM 1 (CER1) |
| AT1G49200 | -5.25656905 | 1.96E-12 | RING/U-box superfamily protein |
| AT5G18404 | -5.19720931 | 1.29E-07 | this gene encodes a small protein and has either evidence of transcription or purifying selection |
| AT5G17220 | -5.12305337 | 0.00165583 | GLUTATHIONE S-TRANSFERASE PHI 12 (GSTF12) |
| AT3G58070 | -4.81136858 | 4.62E-25 | GLABROUS INFLORESCENCE STEMS (GIS) |
| AT2G34925 | -4.80422211 | 1.33E-05 | CLAVATA3/ESR-RELATED 42 (CLE42) |
| AT5G49330 | -4.79177928 | 7.80E-12 | MYB DOMAIN PROTEIN 111 (MYB111) |
| AT2G15020 | -4.7380923 | 3.98E-22 | hypothetical protein |
| AT5G54060 | -4.734528 | 0.04642959 | UDP-GLUCOSE:FLAVONOID 3-O-GLUCOSYLTRANSFERASE (UF3GT) |
