## Supplemental Figures S1-S4 for "The transcriptomic landscape of the photoperiodic stress response in *Arabidopsis thaliana* resembles the response to pathogen infection"

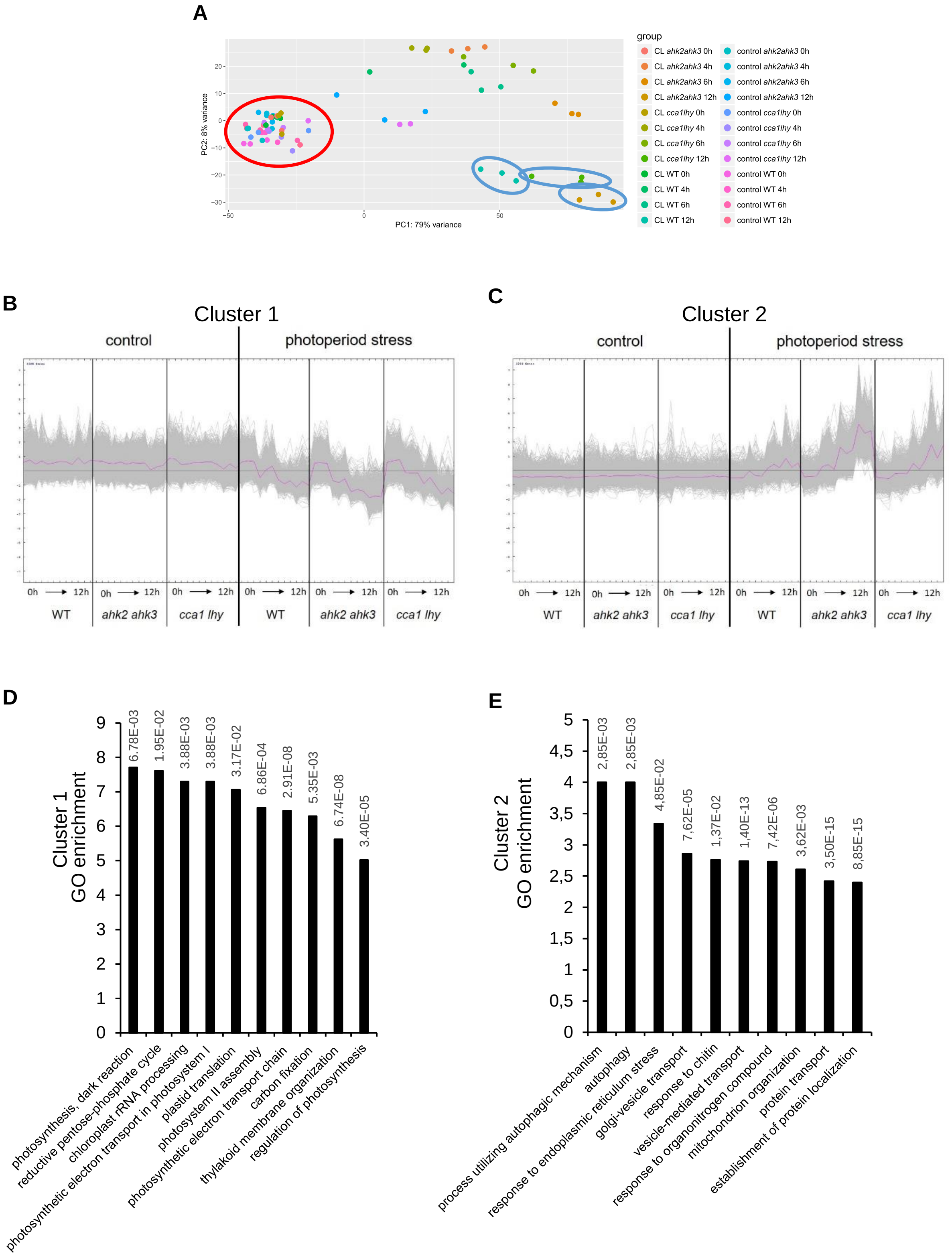


**Supplemental Figure S1. General overview of the RNA-seq dataset.**

(A) PCA of the biological samples. Control samples cluster together (red circle). A strong diversification of light-treated samples is visible with a strong separation of the different genotypes at time point 12 h (blue circles). (B-C) QT clustering of differentially expressed genotype-regulated genes. 52% of all significantly regulated genotype-dependent genes belong to cluster 1 (B) and cluster 2 (C). An overview of the other clusters can be found in Supplemental Figure S2. (D-E) GO term analysis of genes belonging to cluster 1 (D) and to cluster 2 (E).

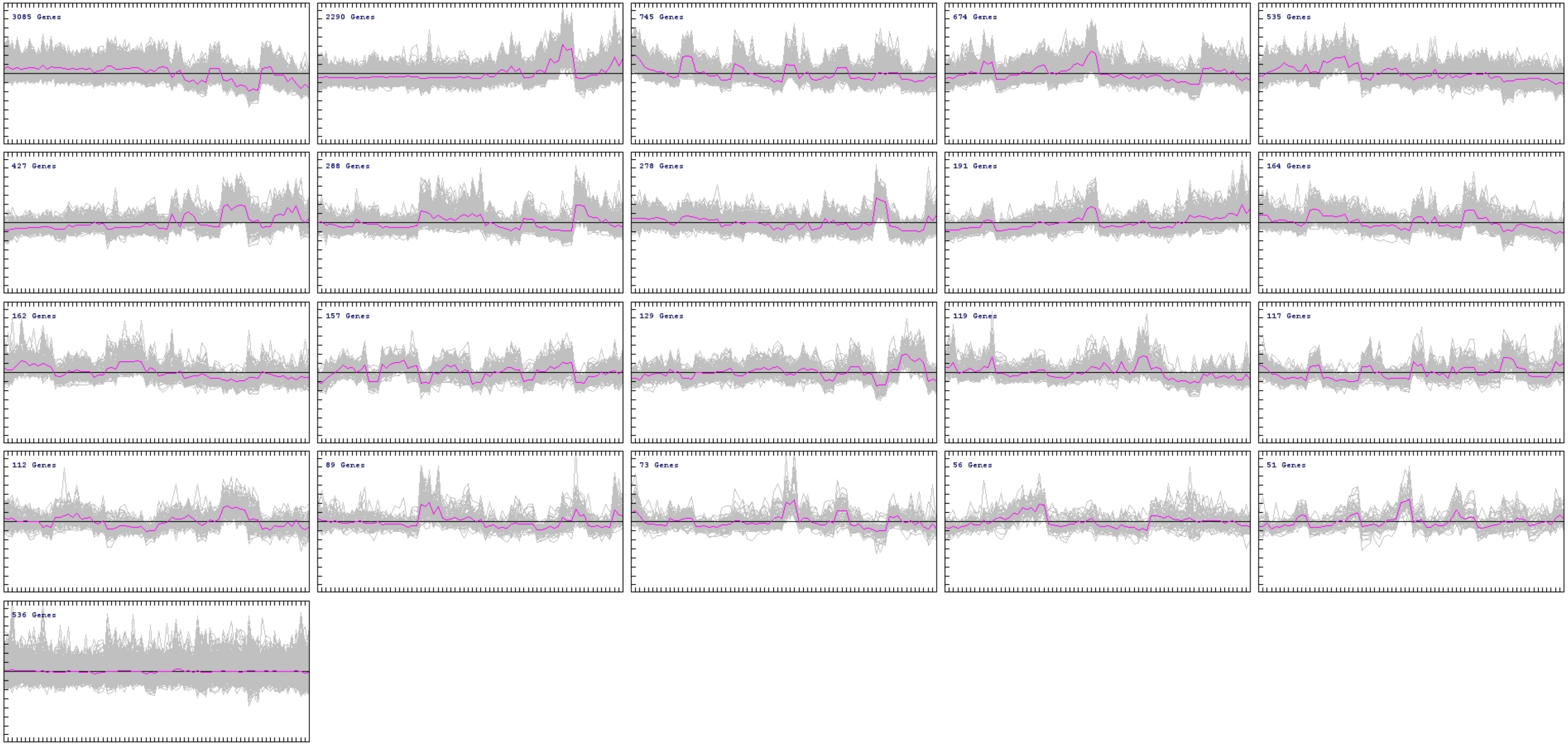


**Supplemental Figure S2. QT clusters of significant genotype-dependent regulated genes.**
QT clustering of differentially expressed genotype-regulated genes.

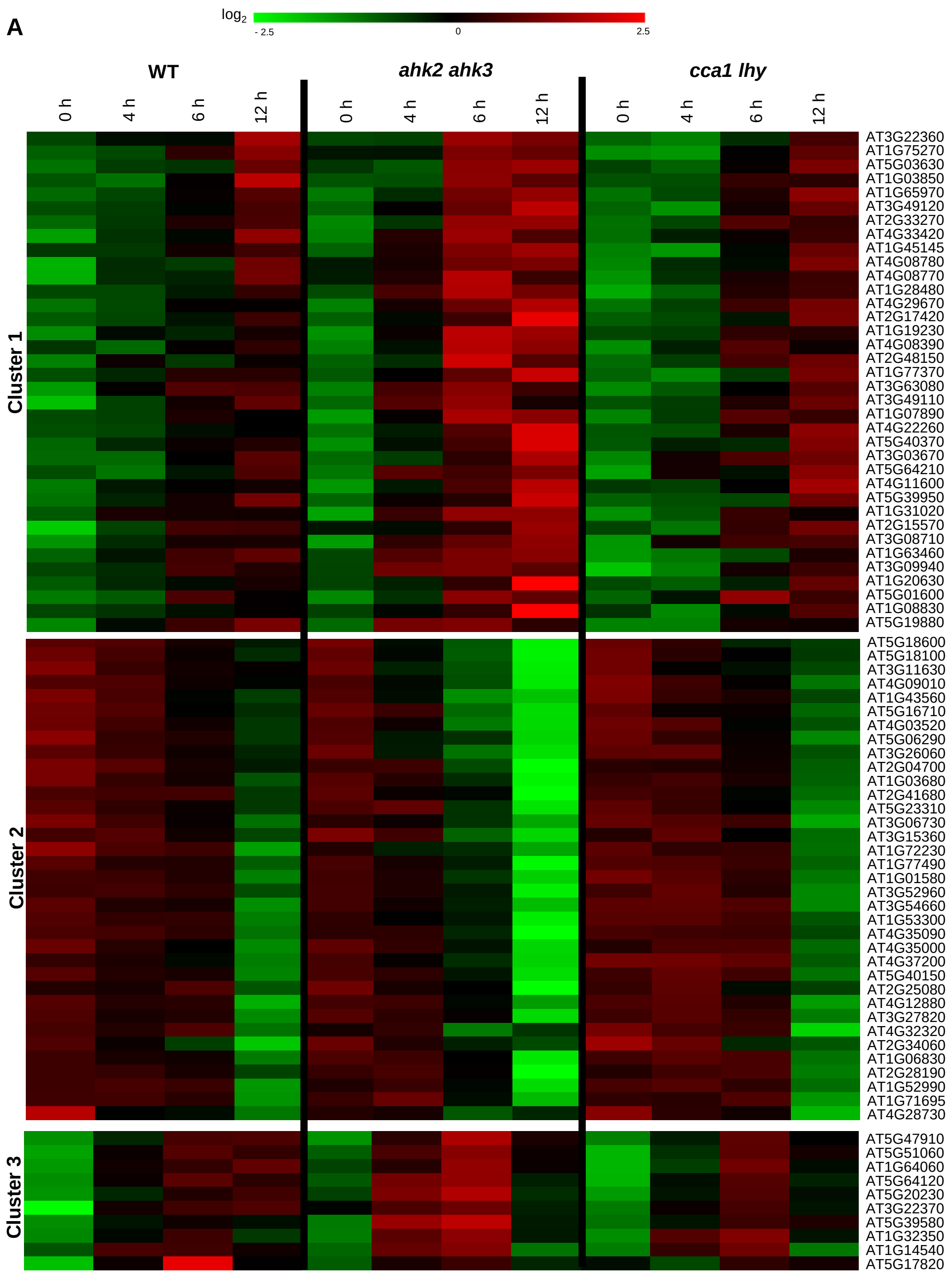


**Supplemental Figure S3. Photoperiod stress causes a strong transcriptional regulation of genes coding for enzymes involved in the scavenging of reactive oxygen species.**
5-weeks-old short day-grown plants were exposed to an extended light period of 32 h followed by a normal short-day night. The experimental setup is shown in Fig. 1A. (A) QT clustering of genes encoding enzymes involved in the scavenging of reactive oxygen species (Pearsons correlation). Three clusters are found: cluster 1 showing an upregulation over time, cluster 2 showing a downregulation over time and cluster 3 showing a strong expression at time points 4 h and 6 h. Depicted are the ratios of PLP versus control plants for each genotype and time point. A complete list of the genes coding for scavenging enzymes used in this analysis including their expression levels is shown in Supplemental Data 3.

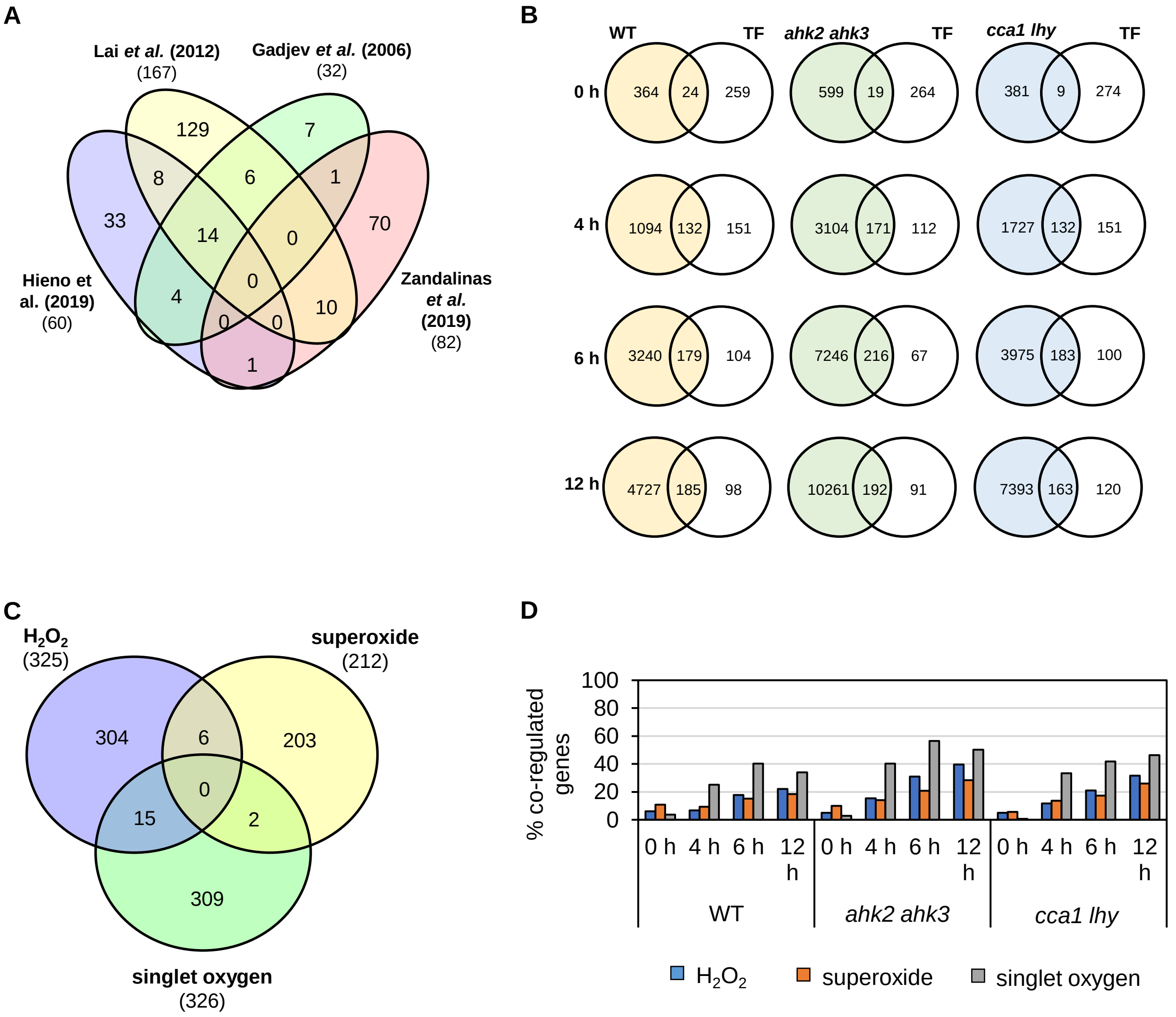


**Supplemental Figure S4. Overlap of photoperiod stress responsive transcription factor genes with different datasets describing ROS-responsive transcription factor genes.**

(A) Venn diagram showing the overlap between the ROS-responsive genes identified by Hieno *et al.* (2019), Zandalinas *et al.* (2019), Lai *et al.* (2012) and Gadjev *et al.* (2006). (B) Venn diagrams showing the proportion of the 283 ROS-responsive genes shown in (A) that are induced or repressed by photoperiod stress in the different genotypes at different time points. (C) Overlap of $H_2O_2$, superoxide and singlet oxygen specific transcript profiles as identified by Gadjev *et al.* (2006). (D) Percentage of DEGs induced by photoperiod stress co-regulated with genes responsive to $H_2O_2$, superoxide and singlet oxygen shown in (C). An overview of the regulation of these transcripts after photoperiod stress is given in Supplemental Data 4 and in Supplemental Data 5.
